# Machine Learning and Directed Evolution of Base Editing Enzymes

**DOI:** 10.1101/2024.05.17.594556

**Authors:** Ramiro M. Perrotta, Svenja Vinke, Raphaël Ferreira, Michaël Moret, Ahmed Mahas, Anush Chiappino-Pepe, Lisa M. Riedmayr, Anna-Thérèse Mehra, Louisa S. Lehmann, George M. Church

## Abstract

As we enter the era of CRISPR medicines, base editors (BEs) emerged as one of the most promising tools to treat genetic associated diseases. However, unintended bystander editing beyond the target nucleotide poses a challenge to their translation into effective therapies. While many efforts have been made in the design of a universal enzyme with minimal bystander editing, the context dependent activity represents a major challenge for base editing-based therapies. In this work, we designed a sequence-specific guide RNA library with 3’-extensions and detected guides that were able to reduce bystander and increase editing efficiency in a context dependent manner. The best candidate was later used for phage assisted non-continuous evolution to find a new generation of precise base editors. Simultaneously, we use protein language models trained on massive protein sequence datasets to find the evolutionarily plausible mutational patterns that can improve deaminase activity and precision. Both strategies provide a collection of precise TadA variants that not only drastically reduced bystander edits, but also was not in detriment of on-target activity. Our findings introduce a guide/enzyme parallel engineering pipeline, which lays the foundation for the development of new personalized genome editing strategies, ultimately enhancing the safety and precision of this groundbreaking technology.

## Main

Base editing has expanded the genome editing toolkit by offering high editing efficiencies, both *in vivo* and *in vitro,* without inducing double-strand breaks (ref). Adenine and cytosine base editors catalyze A•T-to-G•C and C•G-to-T•A transition within a five-nucleotide editing window. When combined with PAMless Cas enzymes, such as SpRY^1,2^, these enzymes have the capacity to induce all four transition mutations and effectively correct the majority of identified human pathogenic Single Nucleotide Polymorphisms (SNPs)^3,4,5^. However, the promiscuous activity of fused deaminases allows the conversion of not only the target nucleotide, but also non-target bases within the editing window. This bystander effect has the potential to result in missense mutations, presenting limitations for the translation into clinical applications^6^.

Despite efforts to minimize bystander edits, they are often proven to be unavoidable, resulting in coding non-silent mutations within the target gene. For adenine base editors (ABEs), assuming the presence of at least one bystander adenine within the activity window, there is a substantial 38% probability that bystander edits will lead to non-silent mutations, potentially with detrimental biological effects^6^. These findings raise profound concerns about the clinical feasibility and safety of base editing technologies.

Recent strategies have been used to decrease bystander activity. For example, through directed mutagenesis of deaminases with narrowed editing windows^7,8,9,10,11^ or the use of gRNAs in combination with SpRY base editors that accommodate the editing window to avoid non-synonymous mutations^11^. Nevertheless, BEs featuring restricted editing windows exhibit diminished editing activity and failed to completely abolish bystander editing^11,12,13^. The BE efficiency and editing pattern are highly influenced by the complex interaction between base editors, gRNAs and target sequences^14^. In addition to the constraints posed by PAM requirements and limited activity windows, BEs exhibits sequence context preferences that significantly impact editing efficiency^6,15^.

Unlocking the full potential of base editing for therapeutic applications requires the development of a robust and adaptable methodology for designing novel base editing strategies that are both efficient and precise. Such an “on-demand” context-dependent approach will ensure efficient and safe customization for a broad range of genetic diseases.

In this work, we first designed and tested a library of 3’-extended sgRNAs, or anchor-guide RNAs (agRNAs), for improving the precision of ABEs. Afterwards, we used the most promising agRNA candidate obtained from the library screening as part of a Phage Assisted Non-Continuous Evolution (PANCE)^16^ system designed to evolve a more precise TadA-8e enzyme. Using a novel dual selection pressure where adaptive mutations by the base editor in the targeted base are necessary, but mutations in bystander adenines are disadvantageous, we discovered several variants with narrowed editing windows and increased activity.

While our use of PANCE successfully generated improved variants, the experimental process itself can be labor intensive and time-consuming. As an alternative, we investigated the potential of leveraging recent progress in deep learning for protein engineering. Specifically, we aimed to design variants with improved fitness in a single step (“one-shot”) without the need for experimental data or manual design effort. We also reasoned that using two orthogonal methods to evolve our protein could lead to the discovery of substitutions that could be mutually beneficial if used together. In line with this, we used machine learning (ML) to computationally design mutants of the TadA-8e enzyme that are more efficient and more precise. Finally, we combined PANCE variants with the ML variants generating a double mutant able to minimize the bystander editing.

The combination of our evolved variants (ABEx’s) with site specific agRNAs represents a tool to design precise base editing systems and thereby to establish a personalized genome editing pipeline for both individual and multiplexed base editing (**Fig.1a**).

**Figure 1.**
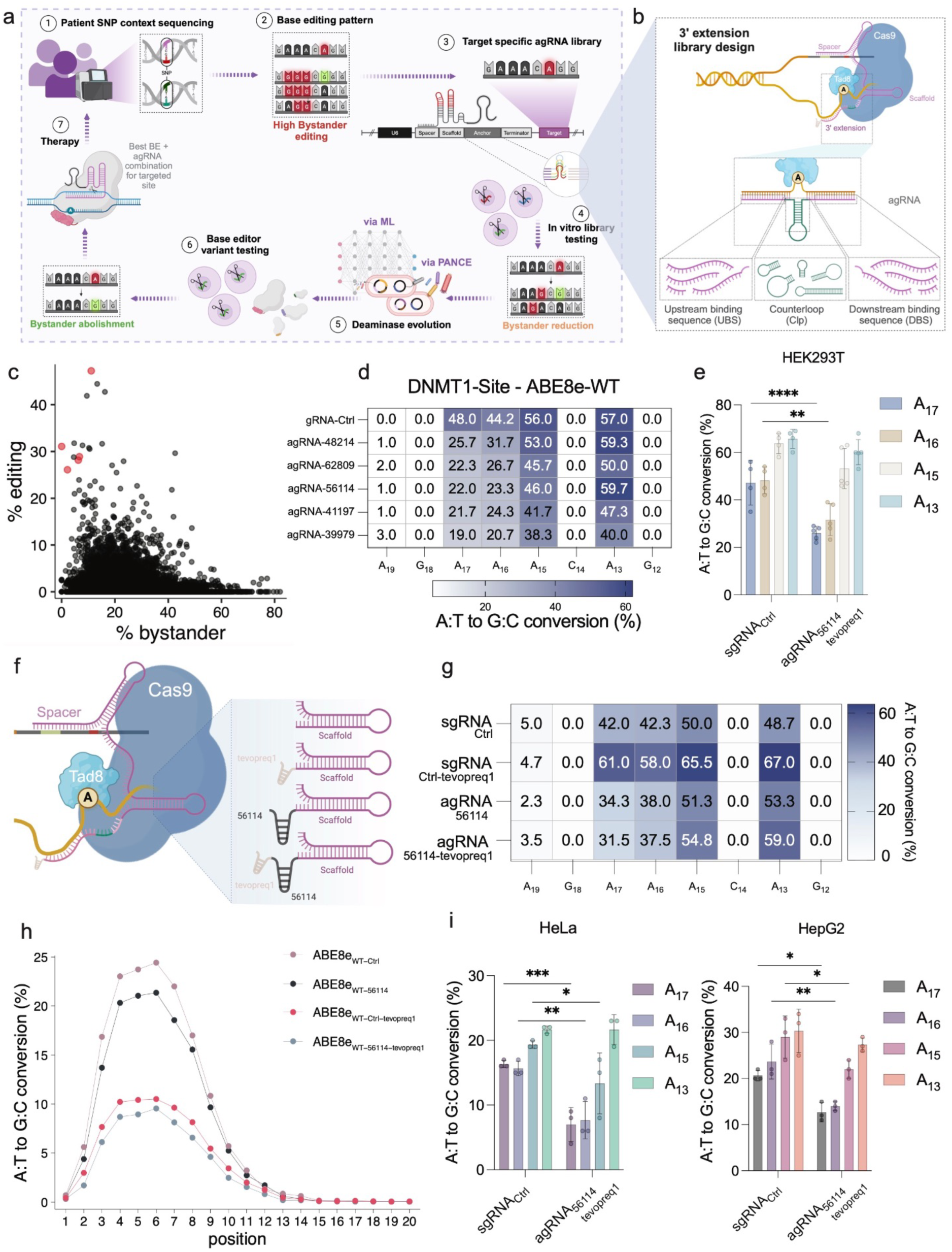
Design and testing of agRNA library and promising candidates: **a,** Schematic workflow of the dual base editor system evolution starting with sequencing the patient specific mutation, testing existing base editing enzymes for that context, and identifying the editing pattern. If there exist bystander mutations, evolving a personalized agRNA for the context, test and optionally evolve novel base editors to generate a “bystander-less” and highly active personalized base editing system for the patient. **b**, Schematic of library design. Library candidates consist of all possible combinations of an array of sequences that bind upstream of the target, hairpin loops, and sequences that bind downstream of the target. **c**, Dot plot representation of the ∼60K agRNA clones’ library after NGS evaluation. In red we highlight agRNA candidates with high efficiency and low bystander editing in the DNMT1 cloned context. **d**, Editing pattern of ABE8e at the human DNMT1 locus in combination with the most promising agRNA candidates. **e**, Editing pattern of ABE8e in combination with agRNA_56114-tevopreq1_ at the human DNMT1 locus in HEK293T cells. **f**, Schematic representation of different agRNA compositions containing different combinations of the anchor and tevopreq1 motif. **g,** Editing pattern of ABE8e and the different guide RNA combinations shown in f at the human DNMT1 locus. **h**, Influence of the agRNA_56114_ with and without tevopreq1 motif on the editing pattern in a ∼12K different pathogenic contexts. **i**, Editing pattern of ABE8e in combination with agRNA_56114-tevopreq1_ at the human DNMT1 locus in HeLa and HepG2 cells. agRNA testing data in the native DNMT1 locus was obtained from n≥3 independent experiments. Data are shown as mean values; error bars, S.D. P value was determined by a two-way ANOVA test (* < 0.05, ** < 0.01, *** < 0.001 and ****, 0.0001).

## agRNAs improves base editing precision

The mechanism underlying the adenosine deamination process involves that the TadA-8e domain engages with the exposed single-stranded region of the PAM-distal nontarget strand (NTS)^17^ (**Supplementary Fig.1a**). The TadA-8e deaminase, when attached to the Cas9 protein, induces specific editing patterns within narrow DNA regions, covering several base pairs. This connection limits the enzyme to act on certain nucleotides, defining what is known as the editing window. Since different DNA contexts are relatively diverse substrates that require the enzyme to accept structure variations within its active site, the DNA strand in the active site has a certain degree of freedom to move in this position.

Our aim was to establish a system that stabilizes the DNA strand within the active site. This restricted movement may result in a smaller editing window thus minimizing the bystander effect. Therefore, we decided to explore the possibility of adding nucleotides to the 3’ end of the gRNA scaffold. These **a**nchor **g**uide **RNA**s (agRNAs) were designed to bind the DNA strand up- and downstream of the DNA region that is later present in the active site of the TadA-8e, thereby stabilizing the loop structure resulting in fewer bases being deaminated (**Fig.1b, Supplementary Fig.1a and Supplementary Table 1-2**).

With this aim, we designed a library of short sequences and entire hairpin structures (counter-loops) at the opposite site of the editing loop to introduce structures that may sterically restrict the movement of the DNA and TadA enzyme even further (**Fig.1b and Supplementary Fig.1a**).

The agRNA library consists of all possible combinations of an array of upstream binding sequences, counter-loops, and downstream binding sequences (**Fig.1b and Supplementary Fig.1a**). Both the up- and downstream sequences bind the DNA strand surrounding the targeted edit. We tested sequences of lengths ranging from 1 to 11 base pair (bp), with all possible start points in a 11 bp window. The counter-loop sequences range from 1 to 33 bp, with the longer ones forming guanine-cytosine (GC)-rich hairpins. This design process yielded an agRNAs library containing ∼60K candidates. To screen the library in a high throughput manner, we constructed a plasmid with the editing target downstream of the agRNA (sensor library). With the agRNA and the target being in a 300-nucleotide window, it is possible to perform next-generation sequencing (NGS) on the sequence in this region after the editing process and analyze the editing pattern for each library clone (**Fig.1a, Supplementary Fig.1a and Supplementary Table 3**).

The tested library was designed to target a site in the human DNMT1 locus, being an optimal candidate for screening, both for high accessibility for editing in HEK293T cells, and the multiple adenines context within the editing window. The ABE8e-spCas9-WT base editor in combination with a non-modified guide (sgRNA_Ctrl_) showed a high editing efficiency for the four adenines in the editing window (A_13_, A_14_, A_15_ and A_17_). Since we detected a slight preference towards the two adenines at position 13 and 15 (**Fig.1d and Supplementary Fig. 1b,d)** with almost the same editing efficiency, we decided to design the library to precisely edit A_13_. With the current editing tools, undesired bystander edits in the selected context remain unavoidable (**Fig.1d and Supplementary Fig.2b**).

We first selected anchors that showed higher efficiency and lower bystander editing in the DNMT1 sensor library (**Fig.1c**). We selected the best 5 candidates to test in the native context in HEK293T cells, and all of them showed a decrease in bystander editing (**Fig.1d**). Clone 56114 was the one that showed higher precision in A_13_ editing, with a significant reduction of 44% in A_17_ and 34% in A_16_ (**Fig.1d and Supplementary Fig.1c-e**). To test the reproducibility of this effect in different cell lines, we also tested the agRNA_56114-tevopreq1_ in Hela and HepG2 cells, obtaining similar bystander reduction patterns (**Fig.1i**).

To determine whether the anchor serves as more than just another 3’ RNA degradation protector like the previously described tevopreq1^18^, we benchmarked various guide combinations with and without the tevopreq1 motif. We observed that the motif increased the editing efficiency of the control guide, probably due to resisting exonucleolytic degradation as has been previously described^18^. The absence of the motif in the agRNA did not affect the bystander reduction effect, suggesting that the reduction is independent of the tevopreq1 motif (**Fig.1g**). To test the ability of influencing the editing pattern in different contexts, we constructed different sgRNA and agRNA libraries targeting ∼12000 pathogenic single nucleotide variants (SNVs) that can be targeted by base editing based on proximity to a PAM^14^. The libraries sgRNA_Ctrl-tevopreq1_ and agRNA_56114-tevopreq1_ drastically decreased the editing efficiency of the ABE8e-spCas9-WT when compared to sgRNA_Ctrl_ (**Fig.1h**). The library with agRNAs_56114_ slightly decreased the editing efficiency. These results suggest that the 3’ modification of the gRNAs maintains the guide’s functionality, but the context influences the efficiency of the anchor and the tevopreq1 motif. Based on this data, we propose that specific agRNAs must be designed for specific contexts to maximize both precision and efficiency.

### Evolution of novel ABEs precise variants via PANCE

Although we observed a robust reduction in bystanders, we were far from generating a perfect edit in position A_13_. To refine the editing, we decided to evolve the enzyme TadA-8e toward decreasing bystander edits and to evaluate how the combination of new variants and agRNA could impact the editing pattern.

Phage-assisted non-continuous evolution has been used in the past to increase the activity of ABEs^19^. Our goal was to design a novel selection method that enables us to evolve variants with decreased bystander edits and that can work with agRNAs. Therefore, we needed a selection pressure that decreases the phage titer in response to bystander edits, but also increases it upon perfect edit, to prevent the evolution of an inactive TadA enzyme. To achieve this, we linked the activity of our base editor encoded on the M13 phage genome with the expression of the *gene III* (encoding for the pIII protein) that is required for phage replication^20^. Since single mutagenesis data for pIII amino acid exchanges are well characterized^21^, we identified groups of 2-3 amino acids that drastically decrease phage replication^21^. Toward this goal, we mutated the codons of selected amino acids in pIII to be T/A rich, whichmake these contexts responsive to A>G editing. We identified three contexts in which bystander edits would result in a decrease in phage titer, while the perfect edit generates an adaptive mutation selecting that also selects for targeted efficiency (**Fig.2a-b**). Each identified pIII variant was cloned into individual plasmids (SP1-3) (**Supplementary Fig.2b**). The selection plasmids also encoded the corresponding agRNA targeting the SNV, as well as a C-terminal Intein-nCas9-SpRY. Once the selection phage carrying the N-terminal Intein-TadA-8e fusion protein infects the cell (**Supplementary Fig.2a**), a functional base editor is expressed, capable of performing the editing (**Fig.2a-b and Supplementary Fig.2c**).

**Figure 2.**
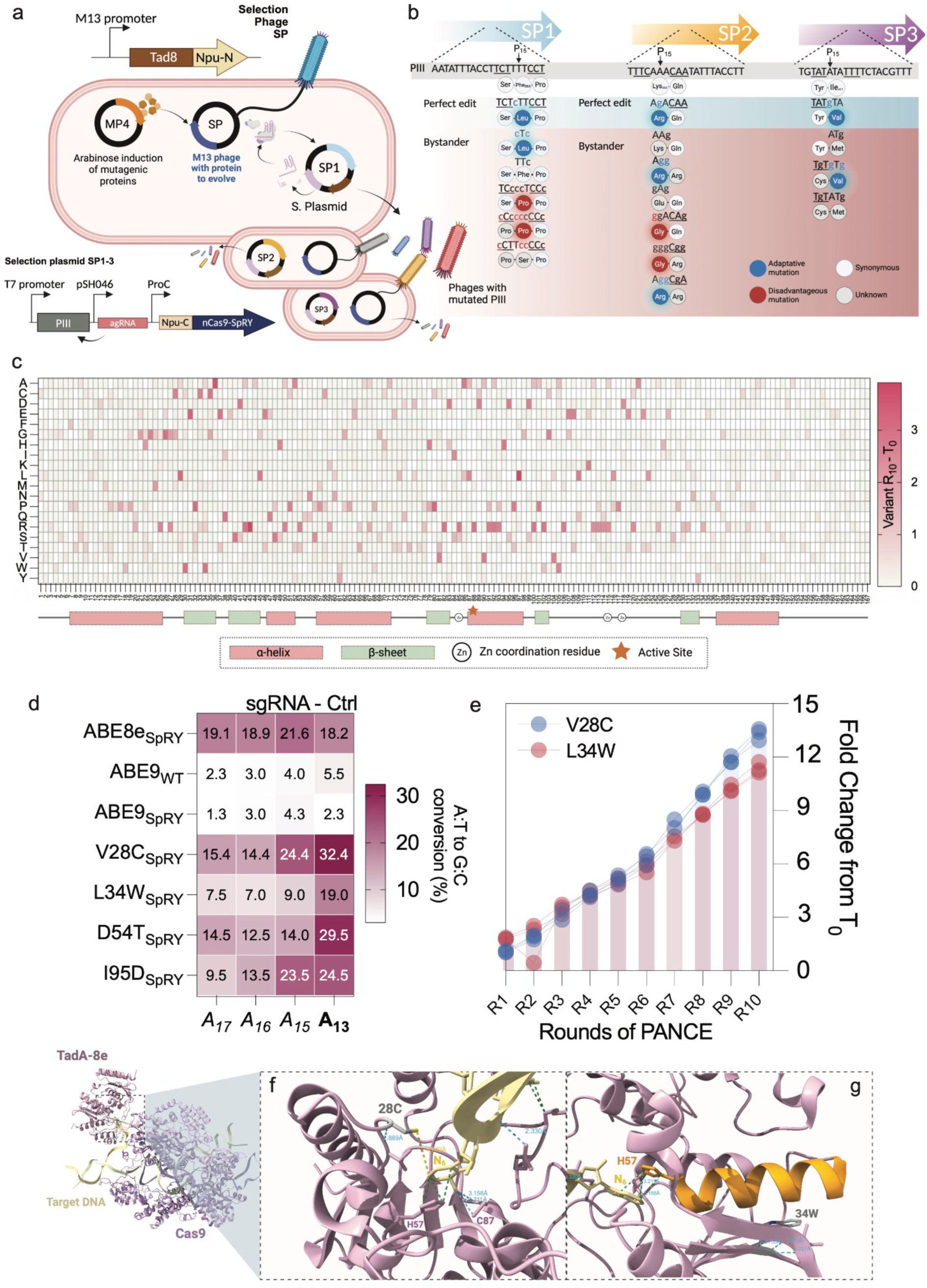
Phage assisted non-continuous evolution of novel adenine base editors. **a**, Schematic representation of the PANCE. The selection phage encoded the TadA-8e adenine deaminase with a C-terminal intein, while the selection plasmids (SP 1-3) the nCas9-SpRY with a N-terminal intein. The three different selection plasmids encode for different pIII sequences carrying a single nucleotide variant (SNV) together with the corresponding agRNA to correct the SNV to the wildtype sequence. **b**, Once the selection phage and the selection plasmid meet in the same cell, the SNVs of SP1-3 can be corrected. If the perfect edit occurs, the sequence is reverted to the wildtype sequence (Blue). The SNVs introduce an amino acid exchange that lowers the phage infectivity (Weiss). Bystander edits can introduce mutations that result in a decrease in the phage replication (Red-Grey). **c**, Mutational landscape after round 10 (R_10_) of evolution. The heatmap represents the mean percentage of amino acid exchanges in R10 minus the amino acid frequencies on the non-evolved phage (n=3 independent experiments) (Supplementary table 5). **d**, Editing pattern of the most promising mutants generated in the PANCE experiment in the human DNMT1 locus in HEK293T cells (n≥3 independent experiments). **e,** Fold change of the amino acid exchanges V28C and L34W representing the two most promising mutants evolved in the PANCE experiment. Dots represent the fold change in the three different replicates of the PANCE evolution. **f-h**, Computational modeling of the structural change resulting from the amino acid exchanges of the V28C and L34W mutants.

Phage DNA was isolated from each PANCE round and sequence using NGS (**Supplementary Fig.2d**). We observed an increase in phage titer after round 4 suggesting an increased activity of the enzyme mutants encoded by the phage (**Supplementary Fig.2e**).

Remarkably the 3 replicates showed almost identical enrichment of the same variant over the course of the evolution (**Supplementary Table 4**). The sequencing data also showed slow enrichment of different mutants with amino acid exchanges, with a maximum enrichment of 3-4% of the new amino acid compared to the wildtype (**Fig.2d**).

Once we determined the TadA mutational landscape, we selected the top 50 most enriched mutations, and individually tested in the DNMT1 site (**Supplementary Fig. 3b,c**). We detected variants with both, higher efficiency and reduced bystander editing pattern at position A_13_, when compared with the Abe8e-SpRY BE (**Fig.2e**). We also benchmarked our PANCE evolved TadA variants against the ABE9 base editor (both WT and SpRY) that showed low editing efficiency and no relative bystander reduction in the DNMT1 site (**Fig.2e and Supplementary Fig.3a-c**). Variants displaying V28C, L34W, D54T, and I95D showed significant potential to generate a perfect edit at position A_13_ (**Fig.2e**). These variants showed an enrichment across the different rounds of evolution suggesting an advantage over other variants in response to the selection pressure (**Fig.2f**). Despite being evolved with the agRNA_56114-tevopreq1_, most of the variants are still functional with the unmodified DNMT1 gRNA (**Supplementary Fig.3b**).

We computationally analyzed the crystal structure of ABE8e to better understand the impact of these mutations on ABE8e. We mutated residues V28C and L34W *in silico*, separately, and compared interactions with surrounding amino acids and nucleotides but no changes in interactions were predicted (**Supplementary Fig.4a-b and SD-1**). We hypothesize that these mutations induce a conformational change in ABE8e that alter interactions of ABE8e residues H57 and C87 with nucleotide 8-Az(26) of the gRNA. Based on the wild-type crystal structure, H57 and C87 are predicted to establish three van der Waals interactions and two hydrogen bonds with 8-Az(26), respectively. In the case of mutant V28C, we hypothesize there could be an approximation of C28 (below the measured distance 5.101Å) to 8-Az(26) of the gRNA from the opposite side than H57 and C87 (**Fig.2f, Supplementary Fig.4b and SD-1**). In line with this, we measured a decrease of 0.77 Å in the distance between residue 28 and 8-Az(26). In the case of mutant L34W, we hypothesize that the tryptophan, which is more hydrophobic than leucine, might alter the orientation of the alpha-helix arm (orange) where residue H57 lies (**Fig.2g, Supplementary Fig.4b and SD-1**).

All in all, non-stop codon based PANCE selection proved to be a powerful tool to evolve base editing mutants that showed decreased bystander editing without losing on-target activity.

### Machine learning guided design of novel precise ABEs

Improving protein fitness traditionally involves screening a vast library of random mutations and selecting those that enhance a desired function under specific pressures (directed evolution). However, our PANCE experiment demonstrates that even with increased fitness, specific mutational patterns across protein families are essential for evolvability. In general, most random mutations will be destabilizing or neutral for protein function (**Fig.3a**).

**Figure 3.**
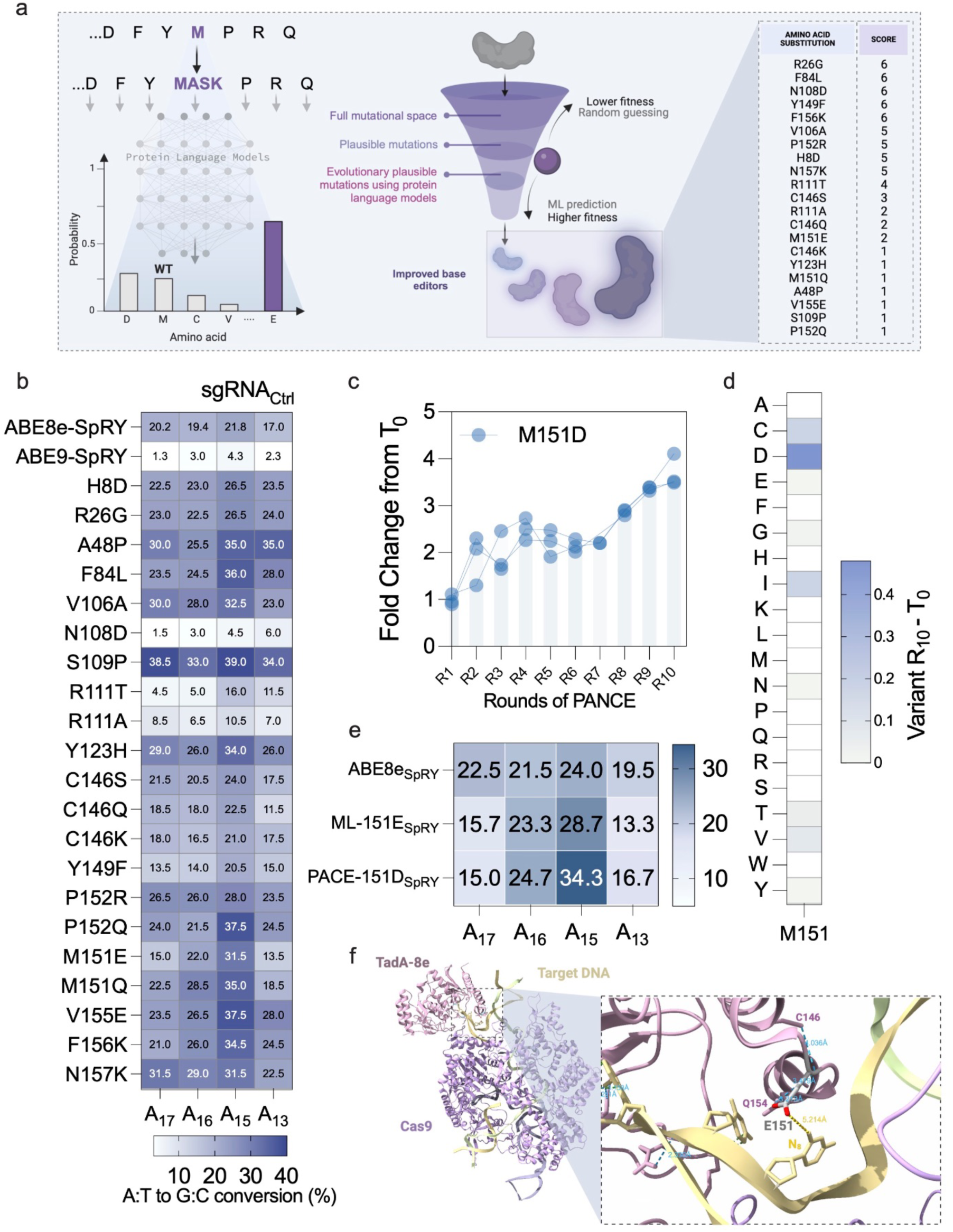
Machine learning guided base editor evolution. **a**, Schematic workflow of the machine learning approach to identify evolutionary plausible mutations (adapted from Hie et al^22)^. **b**, Editing pattern in the human DNMT1 locus caused by ABE single amino acid exchange mutants identified by machine learning algorithm in HEK293T cells (n=3 independent experiments). **c**, Fold change of the amino acid exchange Glutamic Acid (M) to Aspartic Acid (D) at position 151 in the PANCE experiment. **d**, Mean percentage of amino acid exchanges in R10 minus the amino acid frequencies on the non-evolved phage (n=3 independent experiments) of the amino acid exchanges at position 151. **e**, Editing pattern at the human DNMT1 locus of the machine learning identified mutant (M151E) and the PANCE identified mutant (M151D) in HEK2393T cells (n=3 independent experiments). **f**, Computational modeling of the structural change resulting from the amino acid exchanges of the M151E mutant.

In contrast, protein language models trained on massive, non-redundant protein sequence datasets can learn these general, evolutionarily plausible mutational patterns^22,23,24^. This knowledge can be leveraged to predict mutations likely to be beneficial, guiding protein evolution more efficiently. Following training, these models can be used to predict the probability distribution of each amino acid at any given position along a protein sequence, where the probability distribution reflects the knowledge acquired by the models on their training dataset. Positions where the model assigns a higher probability to an amino acid than the wild-type residue are considered more likely than a random pick to yield a positive effect on the protein fitness. This offers a promising and potentially more efficient alternative to traditional directed evolution methods.

In line with this, we used an ensemble of protein language models^22^ to predict mutations likely to yield a positive effect on the TadA-8e sequence, as combining predictions from multiple models has been shown to increase prediction quality, notably on the task of increasing protein fitness^22,23^. We identified 21 evolutionary plausible mutations that we individually tested for their editing pattern in the DNMT1 site (**Fig.3a-b and Supplementary Fig.5a-b**). We observed a different editing pattern, favoring A_15_, in contrast to our PANCE evolved variants where A_15_ showed improved efficiency (**Fig.3b**).

Next, we identified the variant that showed higher A_15_ editing efficiency and reduced A_17/16/13_ bystander edits. We identified M151E as a good candidate for further evaluation. Since this is an evolutionary plausible mutation, we decided to cross-reference with the variants obtained by PANCE. When we evaluated the amino acid substitutions at position 151, we observed an enrichment in aspartic acid across the different rounds of evolution (**Fig.3c-d**). Both glutamic acid and aspartic acid are negatively charged amino acids, suggesting that both independent strategies can identify similar mutations. Both amino acids showed similar editing patterns in the DNMT1 site, with M151D slightly more efficient in position A_15_ (**Fig.3e**).

The computational analysis of ABE8e’s structure highlights a change of interactions for mutant M151E. The wild-type residue M151 forms two hydrogen bonds with C146 and Q154. The mutation E151 allows to form an additional hydrogen bond between the carboxyl group of glutamate (acceptor) and the amino group of Q154 (donor) (**Supplementary Fig.4b**). Glutamate also introduces a negative charge compared to methionine, potentially changing the local conformation, distances (below the measured distance 5.214Å), and interactions between E151 and nucleotide C(25) of the gRNA (**Fig.3f**).

### agRNA and ABEx variants outperform current ABE

After identifying the best agRNA candidate to uncouple undesired bystanders from A_13_ edits at the DNMT1 site, we proceeded to combine agRNA_56114-tevopreq1_ with our most promising ABE variants (**Supplementary Fig.3c and 5b**). We decided to name our most promising variants ABEx’s. ABEx1 (V28C) and 2 (L34W) were generated in the PANCE experiment and ABEx3 (M151E) using ML (**Fig.4a**). ABEx-spCas9-SpRY1 to 3 variants improved efficiency and reduced bystander editing when combined with the agRNA at our target site.

**Figure 4.**
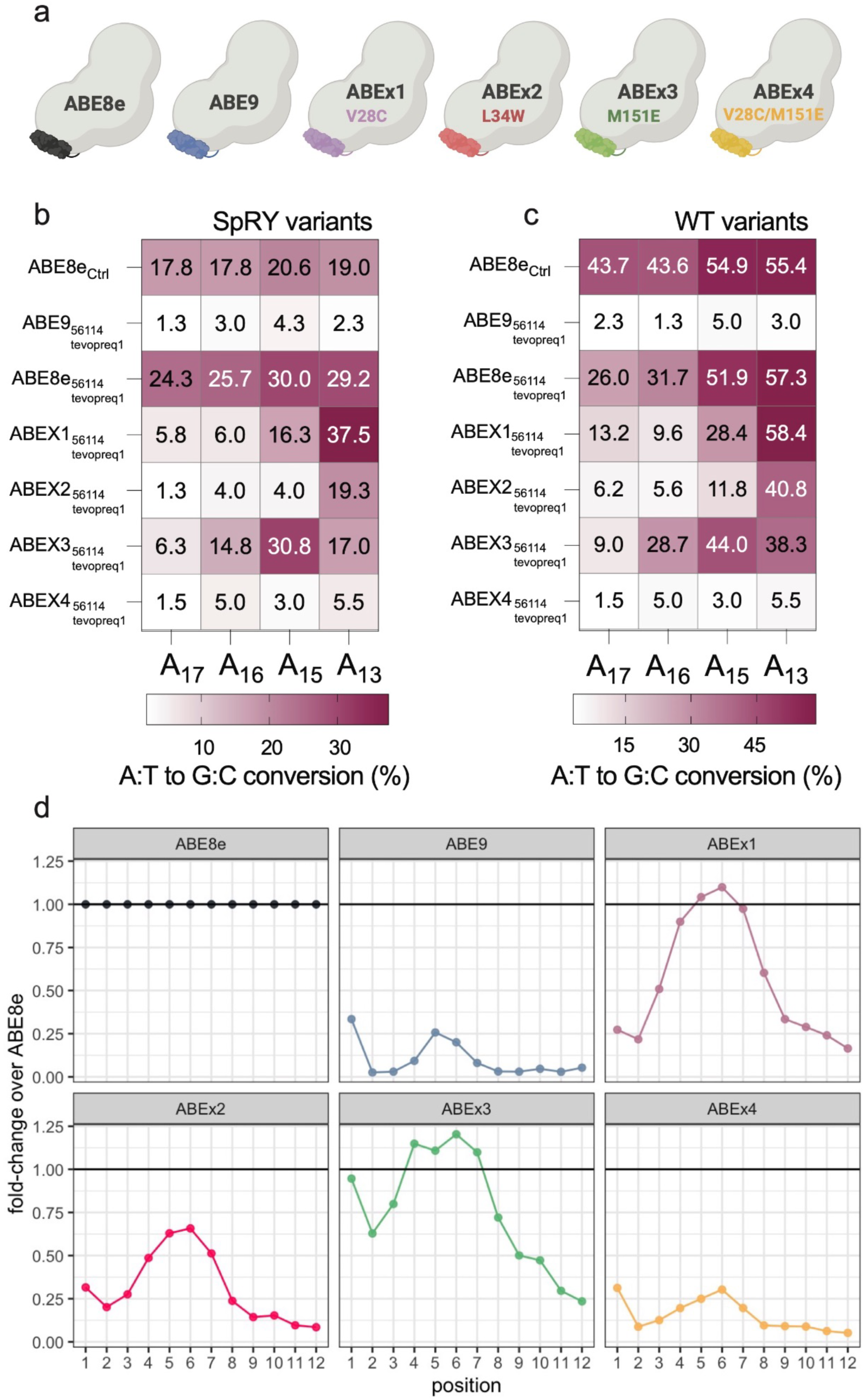
Bystander abolishment by ABEx-agRNA combinations. **a,** ABEx variants and corresponding mutations. ABEx1 and 3 were generated by PANCE, ABEx2 by machine learning and ABEx4 is a combination of both techniques. **b,** Editing pattern in the human DNMT1 locus caused by ABEX spCas9-SpRY variants combined with agRNA_56114-tevopreq1_ in HEK293T cells (n=3 independent experiments). **c,** Editing pattern in the human DNMT1 locus caused by ABEX-spCas9-WT variants combined with agRNA_56114-tevopreq1_ in HEK293T cells (n=3 independent experiments). **d,** ABEX-SpRY variants fold change editing efficiency normalized vs ABE8e, analyzed in ∼12000 different pathogenic contexts.

ABEx1 demonstrated higher editing efficiency at position A_13_ and reduced bystander editing compared to the control, using both sgRNA and agRNA (**Fig.4b and Supplementary Fig.6a-d**). ABEx2 achieved precise and perfect desired editing, exhibiting the same efficiency as ABE8e-SpRY at position A_13_, while also minimizing bystander editing at position A_17/16/15_ (**Fig.4b**). ABEx2 showed precise desired editing. ABEx3, also reduced bystander editing when combined with agRNA_56114-tevopreq1_ and increased efficiency at position A_15_ (**Fig.4b**). We found similar editing patterns when we used the ABEx-spCas9-WT variants (**Fig.4c and Supplementary Fig.6b),** but with higher editing efficiencies. To test a combination of mutations, we created ABEx4 (V28C-M151E). Similar to ABE9, the combination of two mutations on the deaminase seems to abolish the editing activity in this site (**Fig.4b-c**).

However, ABEx4 was functional, even showing higher editing efficiency and bystander reduction, outperforming ABE9 in the HEK site3 (**Supplementary Fig.7a-d**).

A previous strategy to remove bystander editing using the PAMless SpRY variant of the spCas9 was based on the design of guides that move the editing window to isolate the desired base^11^. In line with this, we tested the ability to isolate A_13_ after moving the editing window downstream (-1, -2, -3 bp) and upstream (+1, +2 bp) of our control sgRNA. ABE8e-spCas9-SpRY was not able to isolate A_13_ neither with the sgRNA_Ctrl_ or with agRNA_56114-tevopreq1_. ABEx1, on the contrary, showed increased A_13_ editing with both centered and +1 and +2 sgRNA and agRNA_56114-tevopreq1_ (**Supplementary Fig.8a**). This highlights the importance of targeted design of guides and its further combination with ABEx variants to find the best candidate to fix a particular mutation to maximize efficiency and safety.

ABEx1 with both sgRNA_Ctrl_ and agRNA_56114-tevopreq1_ showed no significant increase in Cas9-dependant (**Supplementary Fig.9a-e**) and independent (**Supplementary Fig.9f-h**) off-target editing, despite the increased on-target activity in position A_13_.

To further investigate our ABEx variants in different contexts, we analyzed the editing pattern using a library with ∼12.000 different pathogenic contexts (NGG, NG-PAMs) (**Fig.4d, Supplementary Fig.10a**)^14^. ABEx1 and ABEx3 exhibited higher editing efficiency than the ABE8e-spCas9-SpRY and narrower editing windows. ABEx2 also reduced bystander editing. However, the L34W mutation impacted the enzyme efficiency. ABEx4 was more efficient than ABE9. However, both double mutants exhibited a big decrease in their efficiency. Additionally, we noticed that ABEx variants have position 15 as the preferred edit while ABE9 optimized position 16 (**Fig.4b, Supplementary Fig.10a and Supplementary Fig.11a**). We also analyzed the effect of the number of As in the editing window, observing a similar editing pattern (**Supplementary Fig.11b-e**).

Overall, we showed that our ABEx mutants can work in a big range of contexts with minimal C to A/G/T editing (**Supplementary Fig.11f**).

The editing fold change, normalized versus the ABE8e-SpRY, revealed that our evolved variants achieved outstanding results, even when compared with variants previously described, like ABE8e and ABE9 (**Fig.4d**). In combination with target specific agRNAs, this parallel evolution base editing pipeline has the potential to revolutionize personalized medicine approaches.

## Discussion

As CRISPR-based therapies enter clinical trials^25,26^, the development of novel strategies to ensure the safety of this technology is crucial. Bystander editing jeopardizes the applicability of this technology, being one of the biggest challenges to ensure a safe and positive outcome for patients, and thus, a widespread use of this revolutionary technology. About 50% precision in half of the pathogenic human SNVs can be achieved by ABEs, but when the editing window contains more than one target base, that number decreases to 26%^14,27^. While engineering the enzyme to obtain base editors with narrower windows seems a streamline solution, previous efforts showed that precision is achieved at the expense of reduction of editing efficiency, imposing a limitation in target sites that can be edited.

In this work, we established a genome editing pipeline that addresses bystander editing challenges via a parallel base editing system evolution. By the design of a 3’ extended gRNAs library, we were able to detect target specific sequences that improved the precision of the ABE8e without compromising efficiency. Moreover, the refinement of agRNA libraries specific for each individual target, creates a novel way to increase the safety of this technology, but also facilitates translational efforts. In parallel, we used a novel PANCE selection system to evolve TadA-8e deaminase variants using the agRNA_56114_. We evolved a battery of enzymes of which most showed bystander reduction in the DNMT1 site and increased efficiency compared to the ABE8e. This proves not only the successful evolution, but also that our PANCE selection system can be used for the evolution of novel precise target-specific base editors.

PANCE evolution paired with NGS sequencing proved to be a powerful tool to identify variants that decrease bystander edits without losing their activity. The evolution pressure in this experiment was relatively low compared to stop-codon based evolution like the PANCE resulting in the identification of ABE8e^19^. PANCE or PACE evolutions with a low selection pressure result in significantly slower enrichment of superior variants, which requires another evaluation than the sequencing of numerous plaques used in most phage assisted evolution experiments^16,19,28,29,30^. The Illumina NGS evaluation has been proven as a good evaluation tool but is only optimal for relatively small target proteins when co-mutations are also supposed to be analyzed, given the current Illumina maximum amplicon size is 600 bp^31^. This highlights the need to establish a workflow using other sequencing techniques like PacBio sequencing for evolution experiments like the one presented in this work but using bigger target proteins.

We also complemented our directed evolution with protein language models finding novel precise variants. A recent work successfully implemented large language models to generate novel gene editors with improved activity compared to spCas9^32^. Our ABEx3 variant showed increased efficiency and reduced bystander editing compared to ABE8e. For the first time, we combined PANCE and machine learning to create a SpRY-ABEx4 enzyme that efficiently reduces bystander editing. Our ABEx’s proteins increased precision and efficiency, creating a more accurate and safer alternative to ABE8e. Our target specific pipeline proposes a straightforward workflow for the design of safe “on demand” base editing therapies. The context dependency of base editing forces us to re-think personalized medicine strategies, especially for rare diseases with low incidence in the population. Our agRNAs create an original alternative to get gene therapy closer to on-demand CRISPR cures.

## Methods

### Bacterial media, reagents and plasmids used in the study

LB and 2xYT media were generated using MP Biomedicals™ media capsules according to the manufacturer’s protocol. For LB and 2xYT agar, 16 g/L agar was added for standard, and 7 g/L agar was added for soft agar. All media was sterilized by autoclaving. Oligos/primers and plasmids used in the study can be found in Supplementary table 6.

### agRNA library generation

The agRNA library consists of an upstream binding sequence (UBS) that is the reverse complement of the downstream sequence of the target, a counter-loop and a downstream binding sequence (DBS) that binds the upstream sequence of the target. The upstream and downstream binding sequences are of different length and have different binding regions in the 1 to 11 bp region upstream and downstream of the target. The counter-loop library consists of 33 different DNA sequences of which the longer one’s form GC rich hairpins. The final library is a library containing every combination of the possible UBS, hairpin, and DBS combinations. A script to generate the hairpin library for a novel context can be found in supplementary material 1.

The agRNA for DNMT1 was ordered as Agilent DNA Oligo Pool (64610 oligos – Supplementary Table 1 and 2). The oligos for the DNMT1 library contained a Gibson overhang, gRNA, gRNA scaffold, agRNA library and a terminator followed by a short DNA sequence used as primer binding site. The target for the DNMT1 library was already cloned on the plasmid used as backbone. The DNA Oligo Pool library was amplified with the oligos Lib_F and Lib_R via PCR. The backbone pU6-tevopreq1-GG-acceptor (Addgene #174038) was PCR amplified using the oligos SplitF and SplitR. The PCR product of the backbone was DpnI digested overnight at 37 °C and both PCR products were purified using the New England Biolabs Monarch PCR & DNA Cleanup Kit according to the Manufacturer’s protocol. The fragments were assembled using the New England Biolabs Gibson Assembly® Master Mix in a 10:1 ratio Library to backbone and 150 ng of the backbone DNA according to the manufacturer’s protocol. 2 µL of the Gibson assembly mix are directly transformed into Lucigen’s Endura Competent Cells and after recovery on 1ml of SOC media, plated on Carbenicillin/Agar plates poured in Nunc™ Square BioAssay Dishes (Cole Palmer #EW-01929-00). For cloning into the backbone LentiGuide-Hygro (Addgene #139462), the library was amplified from the pU6-tevopreq1-GG-acceptor. The backbone was digested using PspXI and Esp3I and cloned by Gibson assembly following the previously described protocol.

For each library at least 10x library coverage of colonies were washed off the plates using LB media and then spun down. The plasmid DNA was extracted using the QIAGEN® Plasmid Plus Midi Kit according to the manufacturer’s protocol from the resulting pellet.

### DNMT1 library testing in HEK cells

HEK293T were purchased from ATCC and maintained in DMEM (Life Technologies) supplemented with 10% FBS (Life Technologies) and kept at 37°C and 5% CO_2_.

20 million cells were seeded in a 225mm^3^ dish and co-transfected the day after using Lipofectamine 3000 (Thermo Fisher) with library plasmid amount corresponding to 1 plasmid per cell and 20 ug of base editor pCMV-T7-ABE8e-nSpCas9-P2A-EGFP (KAC978) (Addgene #185910). Genomic DNA was collected from cells 5 days after transfection.

### Evaluation of the anchor library

Efficiency and precision of the base editor, in combination with the anchor sequences from were evaluated using custom scripts developed in R. The quality of the reads from NGS samples was assessed before further processing. Variant calling techniques were then applied to distinctly identify the perfect edit — conversion of adenine at position 13 to guanine — apart from bystander edits, which encompassed any conversion of the other adenines or combinations involving A_13_. Samples with anchor sequences yielding fewer than 20 reads were excluded to ensure robustness in the data analysis. Furthermore, a quantitative score was devised and calculated using the following formula:

*Score* = (% *Perfect Edit*) / ((% *Perfect Edit* + % *Bystander*)²). Anchors achieving the highest scores and demonstrating at least 20% overall editing efficiency were further characterized experimentally (Supplementary Table 3; Supplementary material 2).

### Context library generation

For the A to G base editor, ∼12000 different gRNAs targeting pathogenic relevant mutations^14^ (Arbab *et al*. 2020) were cloned as a library to test the performance of different hairpins and base editors.

The generation of these context libraries differed from the generation of the agRNA, since extensive recombination events occurred when the gRNA, gRNA scaffold, agRNA and target were introduced as one oligo. To avoid recombination issues, the gRNA and target with 11 bp upstream and 25 bp downstream of the native genomic context were cloned as an oligo lacking the gRNA scaffold and hairpin. Instead of these, the oligo had 2 outward facing BsaI cutting sites with 10 randomized base pairs at that position. The DNA Oligo Pool libraries are amplified with the oligos Lib_F and Lib_R via PCR. The backbone sgBbsI (p2Tol-U6-2xBbsI-sgRNA-HygR) (Addgene #71485) was PCR amplified using the oligos BB_R and BB_F. The PCR product of the backbone was DpnI digested overnight at 37 °C and both PCR products were purified using the New England Biolabs Monarch PCR & DNA Cleanup Kit according to the Manufacturer’s protocol. The fragments were assembled using the New England Biolabs Gibson Assembly® Master Mix in a 10:1 ratio Library to backbone and 150 ng of the backbone DNA according to the manufacturer’s protocol. 2 µL of the Gibson assembly mix are directly transformed into Lucigen’s Endura Competent Cells and after recovery plated on agar plates poured in Nunc™ Square BioAssay Dishes. For each library at least 10x library coverage of colonies were washed off the plates using LB media and then spun down. The plasmid DNA was extracted using the QIAGEN® Plasmid Plus Midi Kit according to the manufacturer’s protocol from the resulting pellet.

The library was then digested using BsaI according to the manufacturer’s protocol and gel purified using the New England Biolabs Monarch Gel Extraction Kit according to the Manufacturer’s protocol. The gRNA scaffold, hairpin and terminator with an inward facing BsaI cutting site up- and downstream were ordered as cloned gene synthesis from IDT. The plasmid was also BsaI digested, and the fragment was purified using the New England Biolabs Monarch Gel Extraction Kit according to the Manufacturer’s protocol. The insert and library were ligated using New England Biolabs T4 DNA Ligase according to the Manufacturer’s protocol with 150 ng backbone and a 10:1 ratio of the insert to the backbone. 2 µL of the ligation mix were directly transformed into Lucigen’s Endura Competent Cells and after recovery plated on agar plates poured in Nunc™ Square BioAssay Dishes. For each library at least 10x library coverage of colonies were washed off the plates using LB media and then spun down. The plasmid DNA was extracted using the QIAGEN® Plasmid Plus Midi Kit according to the manufacturer’s protocol from the resulting pellet.

### Path_Var library testing in HEK cells

HEK293T were purchased from ATCC and maintained in DMEM (Life Technologies) supplemented with 10% FBS (Life Technologies) and kept at 37°C and 5% CO_2_.

For stable Tol2 transposon-mediated library integration, 5 million cells (∼400X coverage) were seeded in 175mm^3^dishes. The following day, cells were co-transfected using Lipofectamine 3000 (Thermo Fisher) with 10ug of Tol2 transposase plasmid (pCMV-Tol2 Addgene # #31823) and 10 ug of Path_Var library. To generate stable library cell lines, cells were selected with hygromycin (25ug/ml) starting the day after transfection and continued for > 2-3 weeks. Following, 10ug of base editor was transfected using Lipofectamine 3000 to 2.5 million cells (∼200X coverage) were seeded the day before in a 100mm^3^ dish. Genomic DNA was collected from cells 5 days after transfection.

### Illumina sequencing and bioinformatic analysis of the libraries

To sequence the libraries before (as quality control) and after testing in the HEK cells, 1ng of the isolated plasmid DNA (QuickExtract DNA Extraction Solution Bioserch Technologies) was amplified using the oligo mix IllSeq_DNMT1_i5_F1-4 and IllSeq_DNMT1_i7_R1-4 using New England Biolabs Q5® High-Fidelity 2X Master Mix according to the manufacturer’s protocol. The forward and reverse oligo contained an i5/i7 overhang for indexing as well as 4-7 Ns to ensure shifting of the sequence to be able to sequence the sequences with high identity. The resulting PCR products were amplified in a second PCR reaction using a compatible combination of the New England Biolabs NEBNext® Multiplex Oligos for Illumina® (96 Unique Dual Index Primer Pairs). The PCR products were purified using the New England Biolabs NEB Monarch® Gel Extraction Kit and quantified via Invitrogen™ Qubit™. A 4 nM Pool of the different libraries was generated and 10 % 4 nM PhiX was added. The Pool was sequenced using the Illumina MiSeq Reagent Kit v2 (300-cycles) according to the manufacturer’s protocol.

### Genome editing of genomic loci

HEK293T, HeLa and HepG2 were purchased from ATCC and maintained in DMEM (Life Technologies) supplemented with 10% FBS (Life Technologies) and kept at 37°C and 5% CO_2_. 20.000 cells were seeded in 96 well plates (Corning) and transfected the day after using jetOPTIMUS® (Polyplus) following manufacturer instructions. 50ng of sgRNA or agRNA (both cloned in pU6-tevopreq1-GG-acceptor) with 150ng of base editor were co-transfected, and cells were harvested after three days for Sanger sequencing (Genewiz) or high throughput sequencing (Quintara Biosciences or in house Illumina miSeq). Spacer sequences of tested loci are found in Supplementary table 7.

### Evaluation of the context library

The evaluation of the context library involved analyzing gRNA libraries, which comprised approximately 12,000 spacer sequences and their respective contextual sequences. The efficiency and editing profiles for each gRNA were established using custom scripts developed in R. First, the target sites—where each spacer binds within the context—were extracted from the NGS reads. Subsequently, for each spacer in the library, all combinations of adenine to guanine conversions were aligned against these extracted sequences. Spacers with fewer than 25 total reads were excluded from the analysis. To quantify overall editing efficiency for the different base editors, the mean A to G conversion rate was calculated by averaging the editing frequencies at each targeted position.

### Generation of the selection phages for PANCE

The PANCE selection phages are carrying the CDS for the ABE8e adenine deaminase instead of the CDS of PIII. The ABE8e adenine deaminase has part of the peptide linker sequence and a C-terminal fused intein CDS to enable it to encode the relatively small protein and not the whole base editor. The phages were generated by PCR amplifying the ABE8e adenine deaminase including the partial sequence of the peptide linker using the oligonucleotides ABE_M13_F and ABE_M13_R. The N-terminal Npu DnaE intein was ordered as gBlock and amplified using the oligonucleotides Npu_ABE_F and Npu_M13_R. The phage backbone was amplified using the oligonucleotides GOI_M13_F and GOI_M13_R using a wildtype M13 phage genomic DNA as a template. All PCRs were performed using NEB Q5® High-Fidelity 2X Master Mix according to the manufacturer’s protocol using 1 ng of template DNA and an annealing temperature of 60 °C. All fragments were digested with DpnI (NEB) overnight at 37°C in the PCR buffer and PCR purified using the NEB Monarch® PCR & DNA Cleanup Kit the next day. The fragments were assembled using the NEB Gibson Assembly® Master Mix according to the manufacturer’s protocol and 3 µL of the reaction were directly transformed into electrocompetent S2060^29^ pJC175e competent cells. The cells were recovered in 500 µL SOC media for 45 min and after that 450 µL and 50 µL were mixed each with 500 µL of freshly grown S2060 pJC175e cells. The cells were immediately mixed with 3 mL soft LB-agar (0.7%) and plated on LB bottom agar plates containing 100 µg/mL carbenicillin. The plates were incubated at 37 °C overnight. Plaques were picked into 50 µL 2xYT media and 1 µL was used as a template for colony PCR using the oligonucleotides ABE_M13_F and Npu_M13_R. Positive phages were amplified by adding the remaining 2xYT media to a freshly grown S2060 pJC175e culture at the OD_600_ of 0.4 and cultivating for 16 h at 37 °C. The cultures were spun down to remove the *E. coli* cells and the phages were precipitated by adding a 20% polyethylene glycol (8000) and 2.5M sodium chloride solution in a 1:4 ratio to the culture supernatant. The mixture is incubated for at least 3h at 4 °C and the phage pellet is resuspended in a PBS buffer, the phage titer was quantified using the Progen Phage Titration ELISA kit and the phages were stored at 4 °C until usage. Additionally, 3 mL of the culture supernatant were used for phage DNA isolation using the Omega Bio-tek E.Z.N.A.® M13 DNA Mini Kit. The isolated DNA was sent to Plasmidsaurus for whole phage DNA sequencing.

### Generation of the selection cells for PANCE

The selection plasmids were designed on the basis of using pJC175e and adding mutations that when edited by the ABE base editor, only perfect edits restore PIII activity while bystander lower pIII activity. The pJC175e backbone was amplified using the oligonucleotides pIII_gBlock_R and pJC175e_Cas_F. The part of the pIII CDS containing the mutation followed by the corresponding guide correcting the introduced mutation downstream as well as the C-terminal DnaE intein necessary to fuse the ABE8e adenine base editor encoded by the phage to the Cas9 encoded by the selection plasmid were ordered as gBlock. Each selection plasmid also encodes the agRNA to fix the mutation on pIII (Supplementary material 3). The three different gBlocks for the three different selection plasmids each encoding a different pIII mutation were amplified via PCR using the oligonucleotides gBlock_R and gBlock_pIII_F. The base Cas9 CDS was amplified from ABE8e plasmid (Addgene #138489) using the oligonucleotides BE_Npu_F and BE_pJC175e_R. All PCRs were performed using NEB Q5® High-Fidelity 2X Master Mix according to the manufacturer’s protocol using 1 ng of template DNA and an annealing temperature of 60 °C. All fragments were digested with DpnI (NEB) overnight at 37 °C in the PCR buffer and PCR purified using the NEB Monarch® PCR & DNA Cleanup Kit the next day. The fragments were assembled using the NEB Gibson Assembly® Master Mix according to the manufacturer’s protocol and transformed into electrocompetent S2060 competent cells. The cells were recovered in 500 µL SOC media for 1h and after that plated on LB agar plates with 100 µg/ML carbenicillin and incubated overnight at 37 °C. Colonies were screened via colony PCr and positive clones were sent to whole plasmid sequencing. The clones with verified sequence were used to generate electrocompetent cells that were then transformed with the mutation plasmid MP4^33^ (Badran). The cells were recovered in 1 mL SOC media and plated on 2xYT agar plates containing 1 % glucose, 100 µg/mL carbenicillin and 25 µg/mL chloramphenicol. 5 colonies were used to start 50 µL shake flask 2xYT 1 % glucose, 100 µg/mL carbenicillin and 25 µg/mL chloramphenicol cultivations. The cultivations were used to freeze 20 % glycerol stocks in 1 mL aliquots after 16 h. Each culture was also used to isolate plasmid DNA for whole plasmid sequencing by Plasmidsaurus to select the glycerol stocks with no mutation in MP4 and the selection plasmid.

### PANCE

The evolution was performed as 10 consecutive batch cultivations in triplicates using a mix of three different selection plasmids in each evolution. For the PANCE experiment,the day prior to the cultivation three 3 mL overnight cultures are prepared using 2xYT media with 1 % glucose, 100 µg/mL carbenicillin and 25 µg/mL chloramphenicol. The cultures are inoculated with the glycerol stock of one of the selection plasmids each. The following day, 3-4 h prior to phage infection 50 mL shake flasks are inoculated with an combined OD_600_ of 0.1 of the pooled overnight cultures with the different selection plasmids. The cells are cultivated in 2xYT with 100 µg/mL carbenicillin and 25 µg/mL chloramphenicol. 30 min prior reaching an OD of 0.4, the cells are induced with 0.5 % arabinose and when the cells reach the OD_600_ of 0.4, the cells are infected with the selection phages at an MOI of 1 for the first selection round. The evolution is performed for 12 h at 37 °C and after that the entire cultivation was spun down and the supernatant was filtered with 0.2 µM filters. For selection round 2-4, 500 µL, for round 5-6 100 µL, and for the remaining rounds 5 µL of the supernatant were used to infect the following evolution. The phage titer after each selection round was determined using the Progen Phage Titration ELISA kit. 3 mL of the each culture supernatant was used for phage DNA isolation using the Omega Bio-tek E.Z.N.A.® M13 DNA Mini Kit. Sequences of ABEx1, 2 and 4 are detailed in Supplementary material 4.

### Illumina sequencing of the PANCE experiment

To sequence the PANCE variants, 1ng of the isolated plasmid phage DNA of each selection round was amplified using the oligo mix IllSeq_ABE_i5_F1-4 and IllSeq_ABE_i5_R1-4 using New England Biolabs Q5® High-Fidelity 2X Master Mix according to the manufacturer’s protocol. The forward and reverse oligo contained an i5/i7 overhang for indexing as well as 4-7 Ns to ensure shifting of the sequence to be able to sequence the sequences with high identity. The resulting PCR products were amplified in a second PCR reaction using a compatible combination of the New England Biolabs NEBNext® Multiplex Oligos for Illumina® (96 Unique Dual Index Primer Pairs). The PCR products were purified using the New England Biolabs NEB Monarch® Gel Extraction Kit and quantified via Invitrogen™ Qubit™. A 4 nM Pool of the different libraries was generated and 10 % 4 nM PhiX was added. The Pool was sequenced using the Illumina MiSeq Reagent Kit v3 (600-cycles) according to the manufacturer’s protocol.

### Evolutionary plausible mutations prediction with machine learning

We followed the approach used by Hie et al., which consists of using an ensemble of six protein language models: ESM-1b^34^ (Rives et al.) and ESM-1v^23^ (Meier et al.), composed itself of five models (accessible at: https://github.com/facebookresearch/esm). Together, the six models are used to predict what amino acids would be more likely than the current wild type amino acid, if any, at each position of the protein sequence given as input. The number of models in the ensemble agreeing on a given prediction (i.e., a specific amino acid substitution) allows to score a given substitution, where a higher score is more likely to yield a positive result.

We applied the ensemble of protein language models to the TadA-8e sequence, which yielded the following predictions (score in parenthesis): R26G (6), F84L (6), N108D (6), Y149F (6), F156K (6), V106A (5), P152R (5), H8D (5), N157K (5), R111T (4), C146S (3), R111A (2), C146Q (2), M151E (2) (Supplementary material 4), C146K (1), Y123H (1), M151Q (1), A48P (1), V155E (1), S109P (1), P152Q (1).

### Computational analysis of structural impact on ABE8e

We used the crystal structure of ABE8e named 6vpc available in the PDB database and the software ChimeraX (version 1.6.1 (2023-05-09)) for visualization and analysis of the structure. We first attempted to predict the structures of relevant mutants in this study using AlphaFold2 (available in the colab notebook from sokrypton). However, no structural change was predicted for these single or double mutants with AlphaFold2, which agrees with previous observations on the limitations of AlphaFold2. In addition, a region relevant to this study involving residues from 151 to 160 of chain F was incorrectly folded by AlphaFold2. Hence, we decided to directly mutate our target residues on the crystal structure of ABE8e, although this assumes there is no conformational or folding change for our mutants.

Using the command line in ChimeraX, we visualized, mutated, and analyzed interactions of target residues. We also analyzed interactions of nucleotides 5-8 in the gRNA with residues of ABE8e. We sought for hydrogen bonds, non-polar (van der Waals) interactions between carbon atoms in the gRNA and protein at a maximum distance of 3.8 Angstroms, and cationic interactions between nitrogen atoms within 5 Å of an aromatic carbon involving our target residues and nucleotides.

## Data availability

All oligonucleotides, sequences of the new Tad8 variants, PANCE sequencing evaluation and gRNAs used in the PANCE selection are included in the Supplementary Data files. Statistic and plots have been generated using PRISM 10 (Graphpad Software LLC). Graphical abstracts were generated with Biorender.com. The ABEx expression plasmids will be deposited on Addgene. All additional plasmids used in this study are freely available from the corresponding authors for academic research use upon request. High-throughput sequencing data of PANCE experiments have been deposited at the NCBI Sequence Read Archive database under accession PRJNA1112370 (to be released)

## Acknowledgments

R.M.P was partially funded by Colossal Biosciences Inc (Award #A47368). S.V was supported by the German Research Foundation (DFG) and S.V, M.M and A.C-P by the US Department of Energy (DOE - Award #DE-FG02-02ER63445). R.F. was supported by the Knut and Alice Wallenberg Foundation (KAW 2019.0581). A.M. was supported by the Ibn Rushd Postdoctoral Fellowship from King Abdullah University of Science and Technology. L.M.R and R.F were supported by the Aging and Longevity-Related Research Fund and EGL Charitable Foundation. We thank the BPF Genomics Core Facility, especially Ashley Ciulla Hurst, at Harvard Medical School for their expertise and instrument availability. We thank Agilent Technologies for the synthesis of oligo pool libraries.

## Author contributions

R.M.P and S.V conceptualized the project, led the analysis, and wrote the manuscript with input from all authors. G.M.C supervised research. R.M.P performed wet lab experiments with help of all authors. R.M.P designed, cloned and tested all libraries. S.V designed the directed evolution systems and libraries. R.F led the bioinformatic analysis. M.M led the machine learning protein design. A.M performed path_var library experiments in HEK293T cells. A.C-P performed the *in-silico* mutagenesis and modeling. L.R developed the evaluation score of agRNA clones. L.R, A.T.M and L.L contributed to wet lab experiments execution.

## Additional information

### Supplementary information

This article contains supplementary material.

**Correspondence and requests for materials** should be addressed to Ramiro Perrotta or George Church.

