## Supplementary information for "Machine Learning and Directed Evolution of Base Editing Enzymes"

Supplementary Figure 1 ……….3
Supplementary Figure 2 ……….4
Supplementary Figure 3 ……….5
Supplementary Figure 4 ……….6
Supplementary Discussion 1 ……..6
Supplementary Figure 5 ……….7
Supplementary Figure 6 ……….8
Supplementary Figure 7 ……….9
Supplementary Figure 8 ……….10
Supplementary Figure 9 ……….11
Supplementary Figure 10 ……...12
Supplementary Figure 11……….13
Supplementary Material 1 ……..14
Supplementary Material 2 ……..15
Supplementary Material 3 ……..14
Supplementary Material 4 ……...…15

**Supplementary Figure 1 | Sanger traces and NGS analysis of ABE8e-spCas9-WT with sgRNA$_{Ctrl}$ and agRNA$_{56114}$. a,** Schematic representation of agRNA library design and workflow. The TadA-8e domain engages with the exposed single-stranded region of the PAM-distal nontarget strand (NTS) fostering deamination. Based on this we deigned a 3' extended agRNA library and cloned into (plasmid). We next transfected 20 million HEK293T cells and analyzed the editing pattern by Illumina sequencing (miSeq V3 300 cycles). **b-c,** Sanger sequencing chromatogram on the DNMT1 genome locus after editing using ABE8e-spCas9-WT with the sgRNA$_{Ctrl}$ and agRNA$_{56114\text{-}tevopreq1}$. d-e, Editing frequencies on the DNMT1 genome locus after editing using ABE8e-spCas9-WT with the sgRNA$_{Ctrl}$ and agRNA$_{56114\text{-}tevopreq1}$ analyzed by NGS.

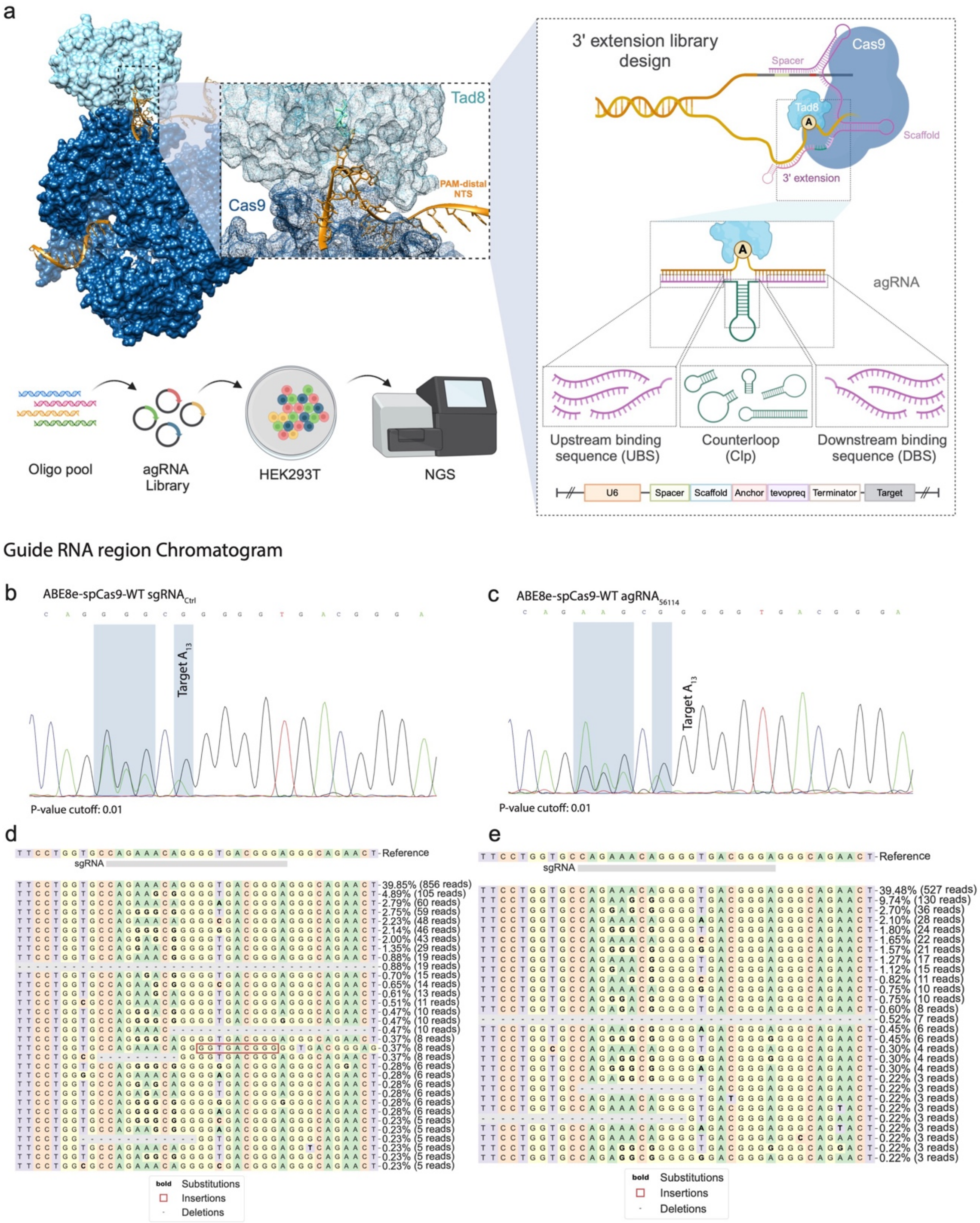

**Supplementary Figure 2 | PANCE constructs and titer.**

**a,** Selection phage design containing the TadA fused with the Npu N-terminal intein. **b,** Schematics of the Selection plasmids containing the gene III, the agRNA and the Cas9 fused with the Npu C-terminal intein. **c,** Mapping of agRNA binding sites on the gene III. **d,** PANCE workflow schematics. **e,** Phage titer across the ten rounds of evolution.

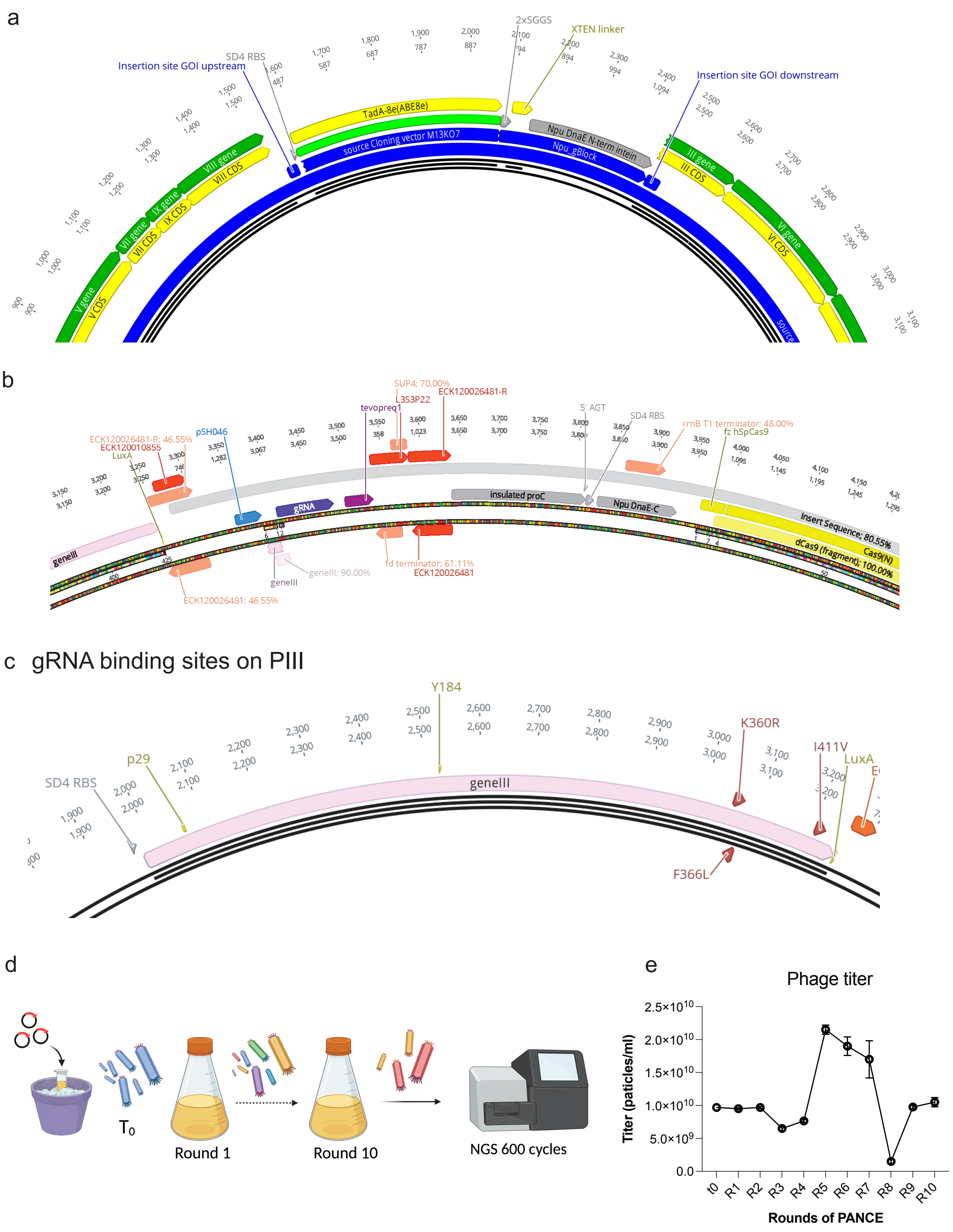

**Supplementary Figure 3 | PANCE variant testing in the DNMT1-Site.**
**a,** RNP editing efficiencies using the ABE8e-SpRY and the ABE9-WT. ABE9 showed low editing efficiency using both plasmids and RNP editing strategies. **b-c,** Editing efficiency of 50 PANCE evolved clones with the $sgRNA_{Ctrl}$ (b) and $agRNA_{56114\text{-}tevopreq1}$.

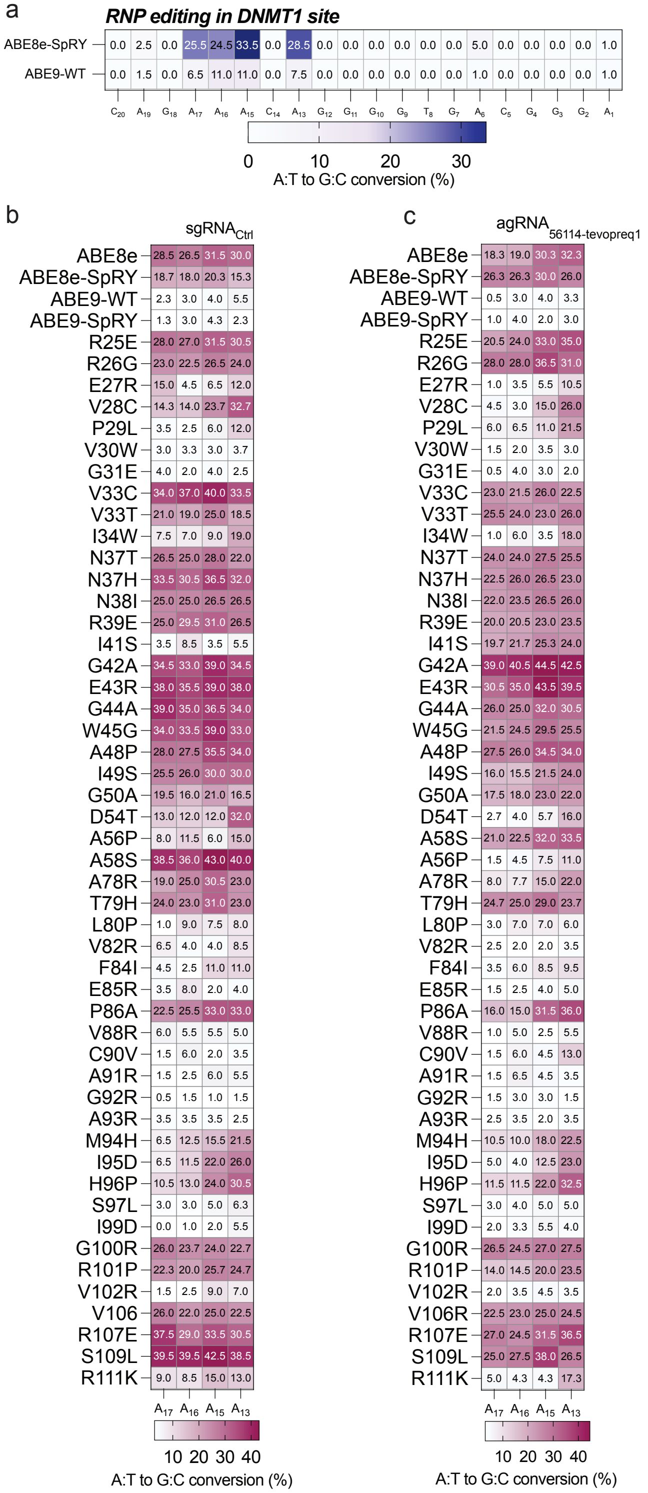

**Supplementary Figure 4 | ABE8e and ABEx variants structure comparison.**

**a,** Structure modeling of ABE8e-WT. Snapshot is at the editing interphase representing the deaminase (pink), the WT mutated aminoacids (grey) and the DNA (yellow). **b,** Number of H bonds formed between amino acids in positions mutated (columns) and neighbor amino acids (rows). Table shows H bonds for those positions with wild-type amino acid (WT), mutated amino acid (mut), and the difference between mutated and wild type (dif).

a

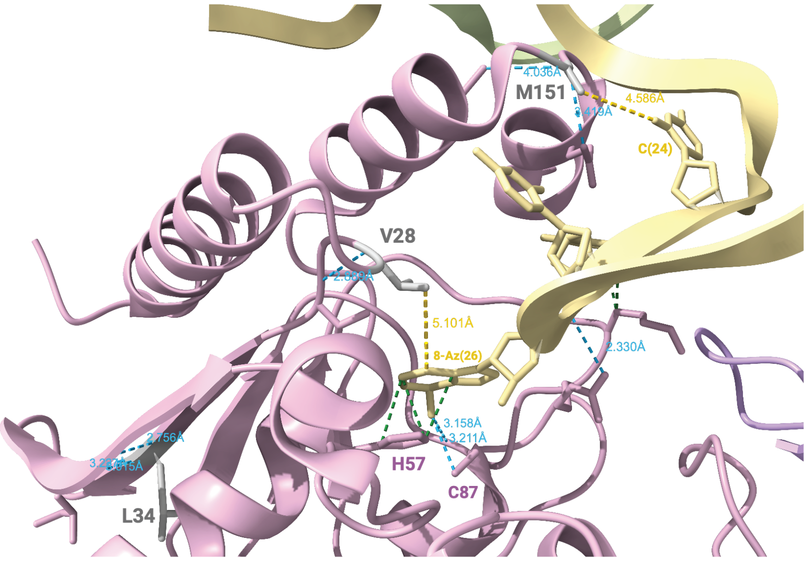


b

| | *V28 (WT)* | *L34 (WT)* | *M151 (WT)* | *C28 (mut)* | *W34 (mut)* | *E151 (mut)* | *V28C (dif)* | *L34W (dif)* | *M151E (dif)* |
|---|---|---|---|---|---|---|---|---|---|
| *V30* | 1 | 0 | 0 | 1 | 0 | 0 | 0 | 0 | 0 |
| *I41* | 0 | 1 | 0 | 0 | 1 | 0 | 0 | 0 | 0 |
| *G42* | 0 | 2 | 0 | 0 | 2 | 0 | 0 | 0 | 0 |
| *C146* | 0 | 0 | 1 | 0 | 0 | 1 | 0 | 0 | 0 |
| *Q154* | 0 | 0 | 1 | 0 | 0 | 2 | 0 | 0 | **1** |

**SD-1.** Both wild-type V28 and mutated C28 are predicted to establish the same interactions with surrounding residues, i.e., a hydrogen bond with V30. However, it is possible that a mutation in residue 28 induces a conformational change that we cannot capture with currently available methods to predict protein structure and it may interact with nucleotide 6 of the gRNA given its proximity (Fig. 2f). Residues L34 and W34 are also predicted to establish the same interactions with surrounding amino acids, i.e., one and two hydrogen bonds with I41 and G42, respectively. Residue 34 is far from the gRNA but the substitution from L to W, which is a more hydrophobic amino acid, might alter the orientation of the alpha-helix arm (orange) where residue H57 lies.

**Supplementary Figure 5 | ML variant testing the DNMT1-Site.**
**a-b,** Editing efficiencies of the 21 ML obtained variants with the $sgRNA_{Ctrl}$ (a) and $agRNA_{56114}$ (b).

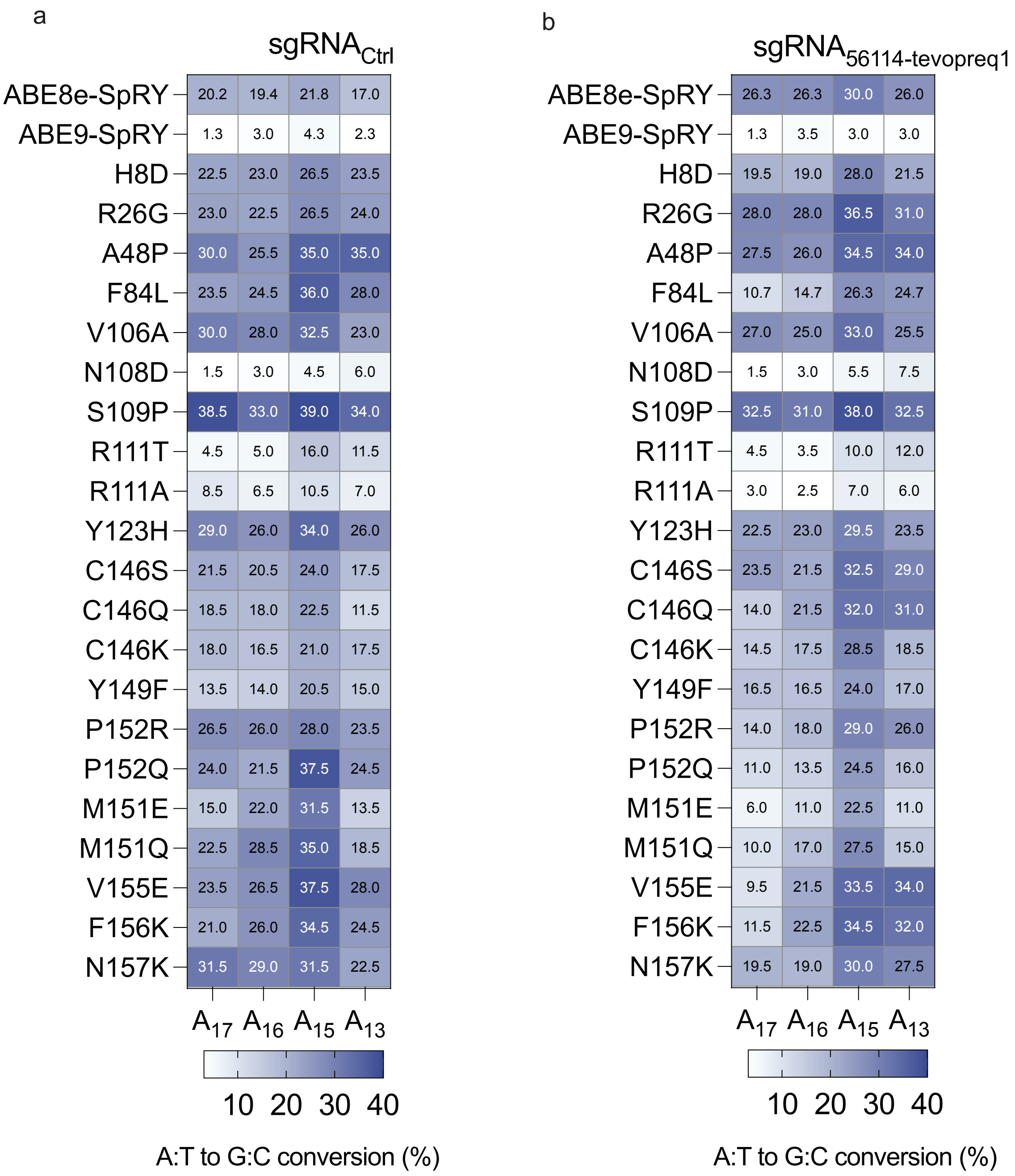

**Supplementary Figure 6 | ABE8e and ABEx1 editing and structure comparison.**
**a-b** Sanger sequencing chromatogram of the DNMT1 genome locus after editing using ABE8e-SpRY with $sgRNA_{Ctrl}$ and $agRNA_{56114}$. b, Mean editing efficiency for WT variants in the DNMT1 locus (n=3 independent experiments).

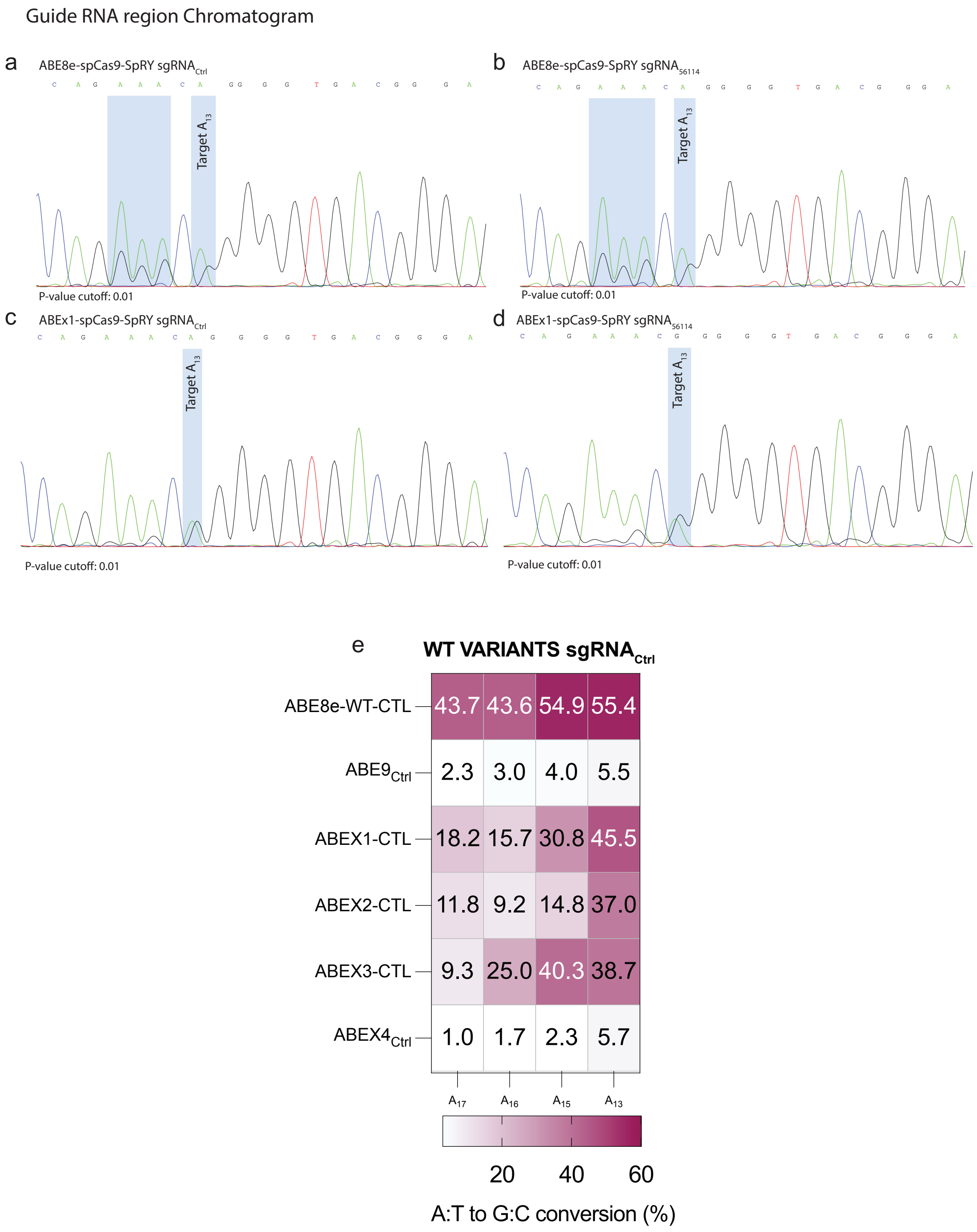

**Supplementary Figure 7 | BE variant testing in HEK-Site3.**
**a,** Editing efficiencies of the HEK site 3 locus using both PANCE and ML variants, together with the double mutants in HEK293T cells. **b-c** Editing efficiencies of the HEK site 3 locus using agRNA$_{56114\text{-tevopreq1}}$ in HepG2 (b) and HeLa (c) cell lines.

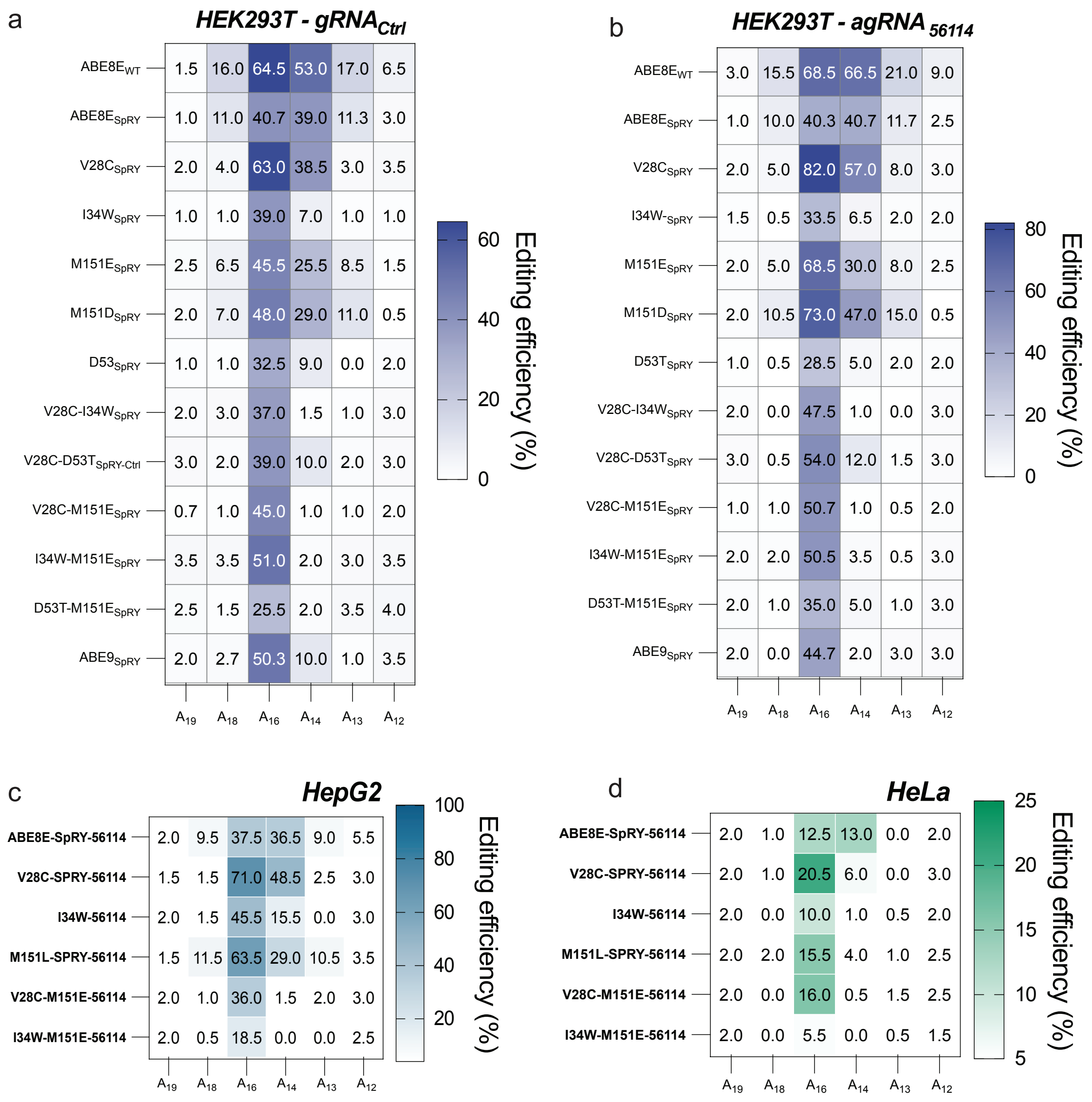

**Supplementary Figure 8 | DNMT1 site editing window movement.**
**a,** Mean editing efficiencies in the DNMT1 locus using different sgRNAs and agRNAs targeting the same site. The different guides are named as -1, -2 or -3 if the editing window is moved upstream and +1 and +2 if downstream the $agRNA_{Ctrl}$ used in this work (centered).

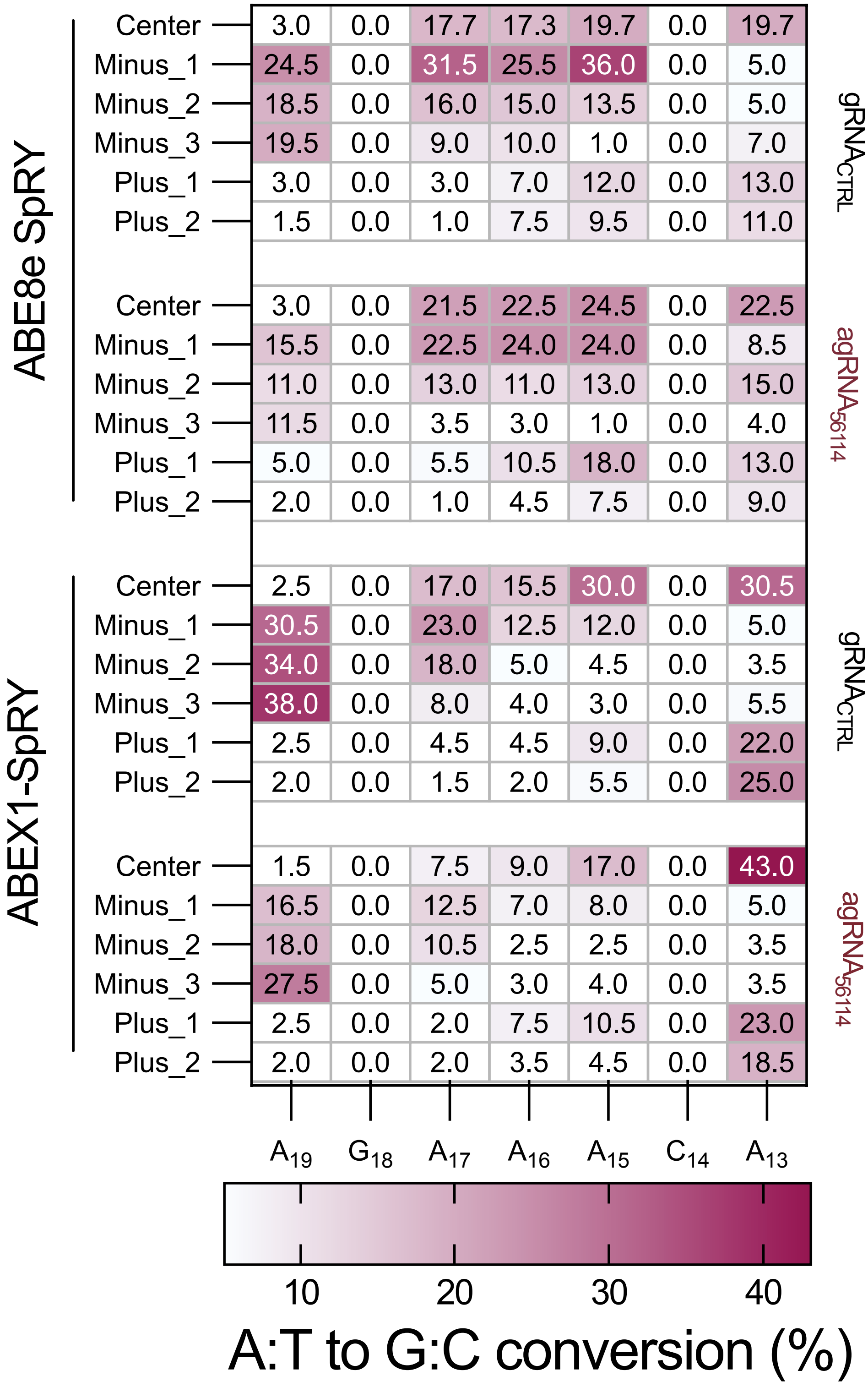

**Supplementary Figure 9 | Cas-dependent and independent off-targets.**
**a,** On-target editing of ABE8e-SpRY and ABEx1-SpRY in the DNMT1 locus. **b-e**, Cas-dependent off-target analysis of ABE8e-SpRY and ABEx1-SpRY in four different loci. **f,** Schematics of the orthogonal R-loop assay. **g-h,** Cas-independent off-target editing of ABE8e-SpRY and ABEx1-SpRY in two different sites.

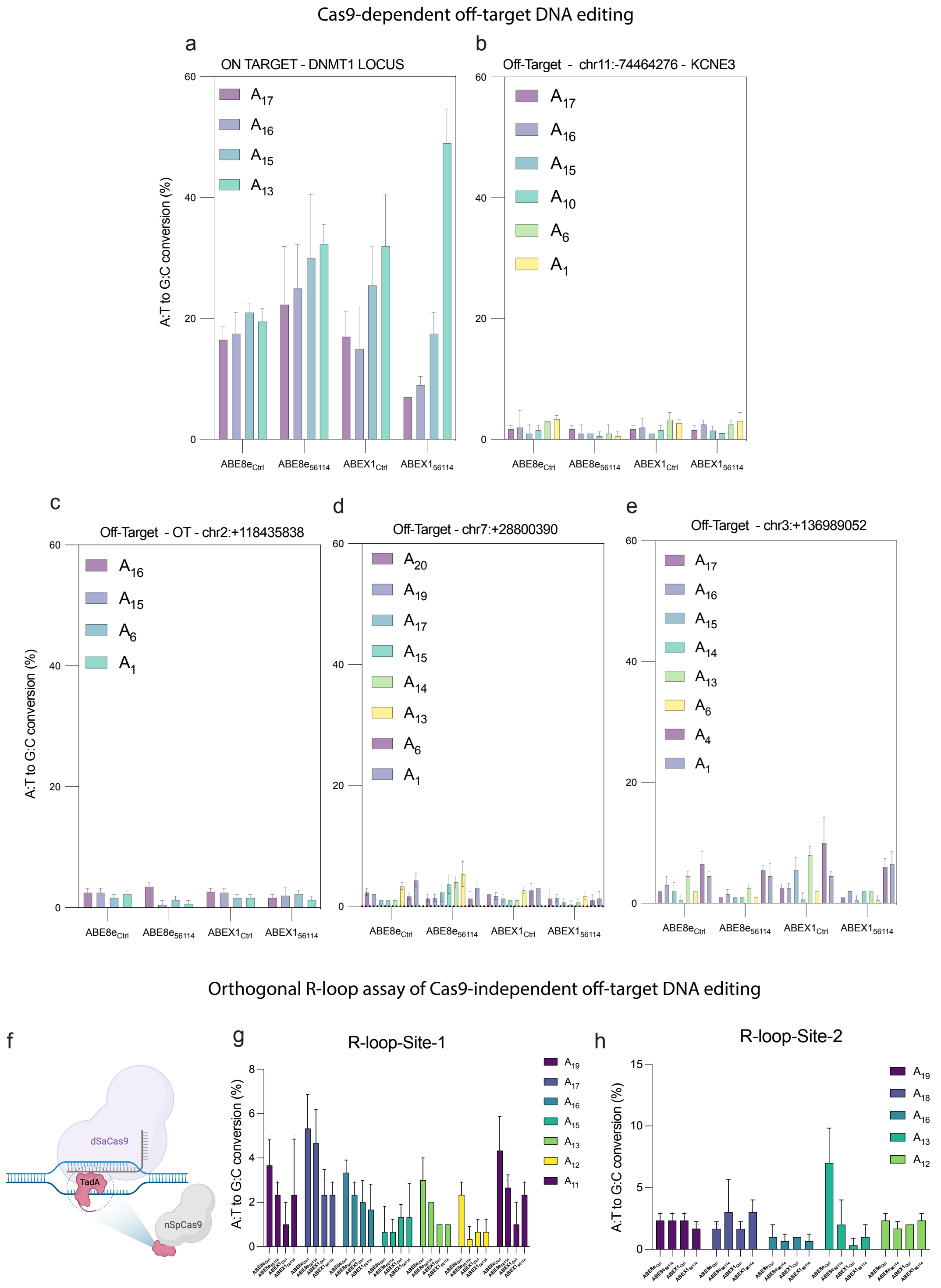

**Supplementary Figure 10 | Path_Var libraries analysis.**
**a,** Editing profile for the different SpRY variants using $sgRNA_{Ctrl}$ and $agRNA_{56114\text{-}tevopreq1}$ compared with ABE8e and ABE9.

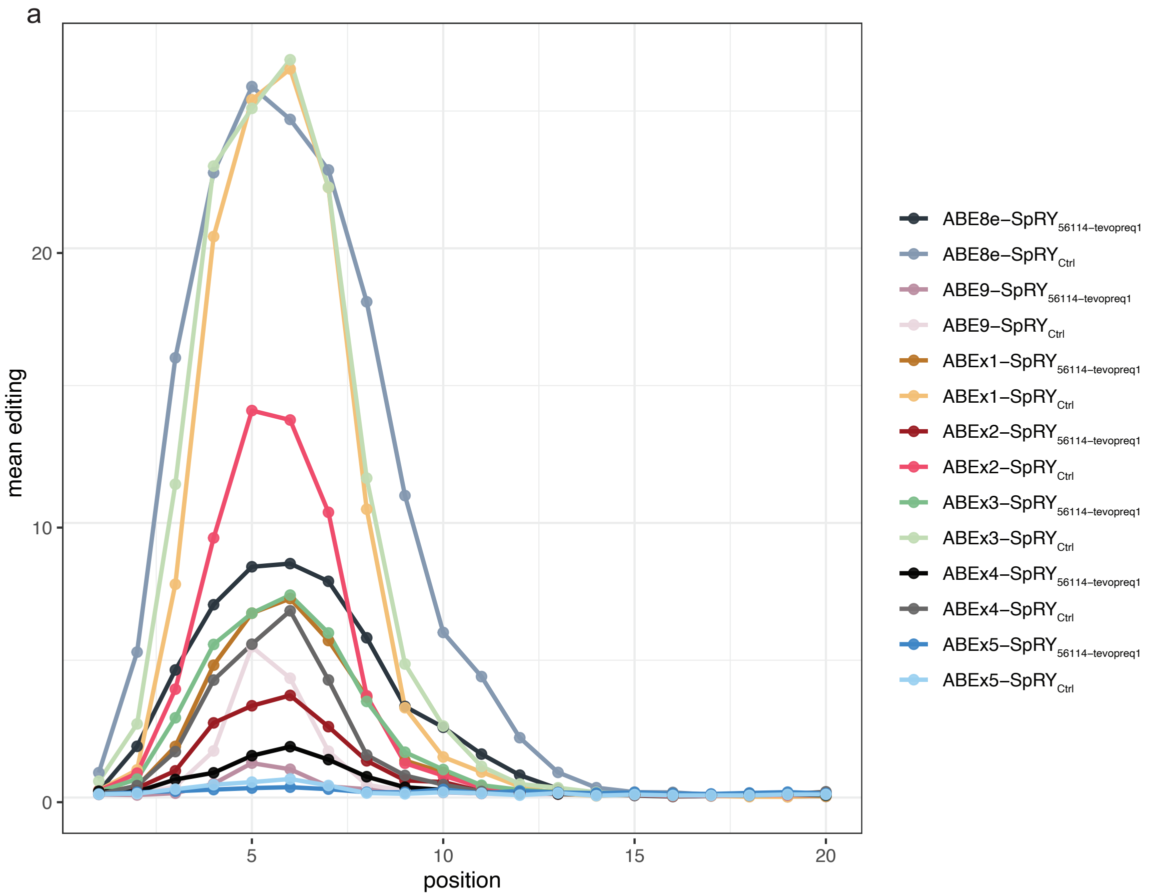

**Supplementary Figure 11 | Path_Var libraries analysis.**
**a,** Read counts of each sample after Illumina sequencing. **b,** Example of the quality score of read1 and read 2. **c,** Editing profile for the different SpRY variants using $sgRNA_{Ctrl}$ compared with ABE8e and ABE9. **d-g,** Editing profile for the different SpRY variants using $sgRNA_{Ctrl}$ compared with ABE8e and ABE9 when more than 2 As are present in the editing window. **h,** C to A, C to G and C to T editing profiles using $sgRNA_{Ctrl}$ for the different SpRY variants compared with ABE8e and ABE9

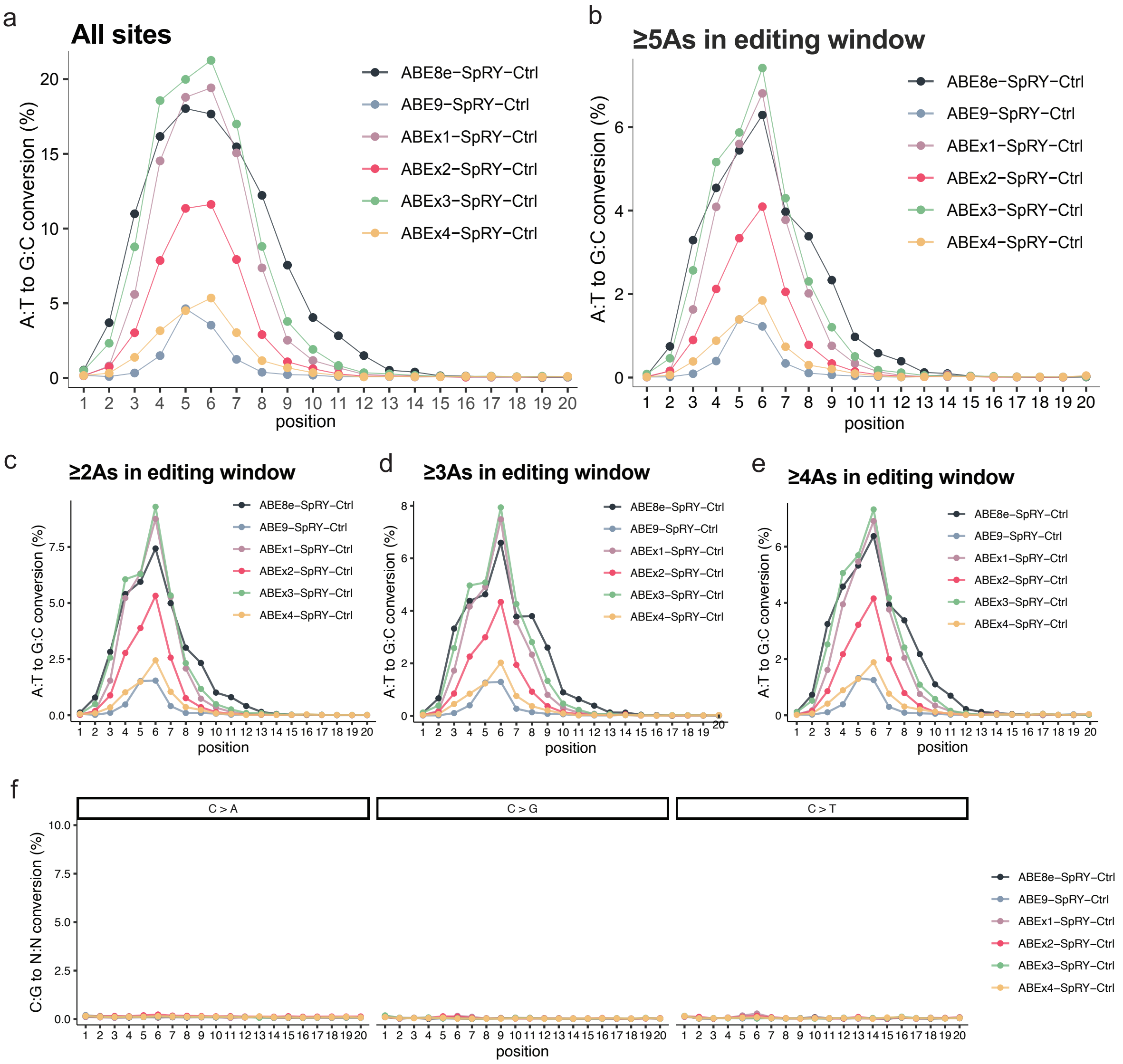

**Supplementary Material 1.** Python script for the generation of agRNA libraries for custom contexts.

import openpyxl
input_sequence = "TACCAGGACCCGCTCAATGTC" loop_sequences = ["GCGCGGCTTCGCGC", "GCGCGGCTCGCGC", "GCGCGCTTCGCGC", "GCGCGTTCGCGC", "GCGCGTCGCGC", "CGCGGCTTCGCG", "CGCGCGCGGCTCGCG", "CGCGCTTCGCG", "CGCGTTCGCG", "CGCGTCGCG", "GCGGCTTCGC", "GCGGCTCGC", "GCGCTTCGC", "GCGTTCGC", "GCGTCGC", "CGGCTTCG", "CGGCTCG", "CGCTTCG", "CGTTCG", "CGTCG", "GGCTTC", "GGCTC", "GCTTC", "GTTC", "GTC", "CCCC", "GGGG", "CCC", "GG", "CC", "GG", "C", "G"]

**# Reverse complements the input sequence**
complement = {'A': 'T', 'T': 'A', 'C': 'G', 'G': 'C'} reverse_complement = ".join([complement[base] for base in input_sequence[::-1]]) UBS = set() DBS = set() # Extract all possible base sequences from the first 10 bases of the reverse complement of the input sequence, and add them to the UBS setstick for i in range(10): for j in range(i+1, 11): UBS.add(reverse_complement[i:j])

**# Extract all possible base sequences from the last 11 bases of the reverse complement of the input sequence, and add them to the DBS set for i in range(11, 21):**
for j in range(i+1, 22): DBS.add(reverse_complement[i:j]) # Convert the UBS and DBS sets to lists, and remove any duplicatesUBS = list(set(UBS)) DBS = list(set(DBS)) # Create a list to store all possible combinations of UBS+Loop+DBS combinations = []

**# Iterate over each UBS sequence, loop sequence, and DBS sequence, and combine them to form a new sequence for u in UBS:**
for l in loop_sequences: for d in DBS: new_sequence = u + l + d combinations.append(new_sequence)

**# Write the combinations to an Excel file**
workbook = openpyxl.Workbook() worksheet = workbook.active for combination in combinations: worksheet.append([combination]) workbook.save("combinations.xlsx") print("Unique Base Sequences (UBS):") print(UBS) print("\n") print("Unique Downstream Base Sequences (DBS):") print(DBS)print("\n") print("All possible combinations of UBS+Loop+DBS:") print(combinations)

**Supplementary material 2:** Anchor sequences in Figure 1d.

**LIB_48214** ACCGCGCTTCGCGTGGCACCA
**LIB_62809** CTCGCGGCTTCGCGTGGCAC
**LIB_56114** CGCGCGTTCGCGCGG
**LIB_41197** CACGCGGCTTCGCGGGCACCA
**LIB_39979** CACGCGCGTTCGCGCTGGCACCA

**Supplementary material 3:** Selection plasmid agRNA design.

pSH046-Spacer-Scaffold-anchor$_{56114}$-Tevopreq1

SP1 (F366L)
TTGACAGGTGAACGCTCAGCTCTTATAATGCCTATAGGAAAAGAAGGTAAATATTGT
TTTAGAGCTAGAAATAGCAAGTTAAAATAAGGCTAGTCCGTTATCAACTTGAAAAA
GTGGCACCGAGTCGGTGCCGCGCGTTCGCGCGGCGCGGTTCTATCTAGTTACGCGTT
AAACCAACTAGAA

SP2 (K360R)
TTGACAGGTGAACGCTCAGCTCTTATAATGCCTATTTTCAAACAATATTTACCTTGTT
TTAGAGCTAGAAATAGCAAGTTAAAATAAGGCTAGTCCGTTATCAACTTGAAAAAG
TGGCACCGAGTCGGTGCCGCGCGTTCGCGCGGCGCGGTTCTATCTAGTTACGCGTTA
AACCAACTAGAA

SP3 (I411V)
TTGACAGGTGAACGCTCAGCTCTTATAATGCCTATTGTATATATTTTCTACGTTTGTT
TTAGAGCTAGAAATAGCAAGTTAAAATAAGGCTAGTCCGTTATCAACTTGAAAAAG
TGGCACCGAGTCGGTGCCGCGCGTTCGCGCGGCGCGGTTCTATCTAGTTACGCGTTA
AACCAACTAGAA

**Supplementary material 4:** Tad8 variant sequences from Figure 4b and c**.**

BPNLS-Tad8

ABEx1 (V28**C**)
MKRTADGSEFESPKKKRKVSEVEFSHEYWMRHALTLAKRARDERE**C**PVGAVLVLNNR
VIGEGWNRAIGLHDPTAHAEIMALRQGGLVMQNYRLIDATLYVTFEPCVMCAGAMIHS
RIGRVVFGVRNSKRGAAGSLMNVLNYPGMNHRVEITEGILADECAALLCDFYRMPRQV
FNAQKKAQSSIN

ABEx2 (L34**W**)
MKRTADGSEFESPKKKRKVSEVEFSHEYWMRHALTLAKRARDEREVPVGAV**W**VLNNR
VIGEGWNRAIGLHDPTAHAEIMALRQGGLVMQNYRLIDATLYVTFEPCVMCAGAMIHS
RIGRVVFGVRNSKRGAAGSLMNVLNYPGMNHRVEITEGILADECAALLCDFYRMPRQV
FNAQKKAQSSIN

ABEx3 (M151**E**)
MKRTADGSEFESPKKKRKVSEVEFSHEYWMRHALTLAKRARDEREVPVGAVLVLNNR
VIGEGWNRAIGLHDPTAHAEIMALRQGGLVMQNYRLIDATLYVTFEPCVMCAGAMIHS
RIGRVVFGVRNSKRGAAGSLMNVLNYPGMNHRVEITEGILADECAALLCDFYR**E**PRQV
FNAQKKAQSSIN

ABEx4 (V28**C**/M151**E**)

MKRTADGSEFESPKKKRKVSEVEFSHEYWMRHALTLAKRARDERE**C**PVGAVLVLNNR
VIGEGWNRAIGLHDPTAHAEIMALRQGGLVMQNYRLIDATLYVTFEPCVMCAGAMIHS
RIGRVVFGVRNSKRGAAGSLMNVLNYPGMNHRVEITEGILADECAALLCDFYR**E**PRQV
FNAQKKAQSSIN
